# Transposable elements contribute to the evolution of host shift-related genes in cactophilic *Drosophila* species

**DOI:** 10.1101/2024.03.27.587021

**Authors:** D. S. Oliveira, A. Larue, W. V. B. Nunes, F. Sabot, A. Bodelón, M. P. García Guerreiro, C. Vieira, C. M. A. Carareto

**Author notes:** Daniel Siqueira de Oliveira. Anaïs Larue. François Sabot. Alejandra Bodelón. María Pilar García Guerreiro.

## Abstract

Host shifts in insects have been considered a key process with the potential to contribute to reproductive isolation and speciation. Both genomics and transcriptomics variation have been attributed to such a process, in which gene families with functions associated with host localization, acceptance, and usage have been proposed to evolve. In this context, cactophilic *Drosophila* species are an excellent model to study host shift evolution, since they use a wide range of cacti as hosts, and many species have different preferences. Transposable elements are engines of genetic novelty between populations and species, driving rapid adaptive evolution. However, the extent of TEs’ contribution to host shift remains unexplored. We performed genomic and transcriptomic analyses in six genomes of cactophilic species/subspecies to investigate how TEs interact with genes associated with host shift. Our results revealed enrichment of TEs at promoter regions of host shift-related genes, with ∼39% of the odorant receptors containing their transcription factor binding sites within TEs. We observed that ∼50% of these TEs are *Helitrons*, demonstrating an unprecedented putative *cis*-regulatory role of *Helitrons* in *Drosophila*. Differential expression analysis between species with different preferred hosts revealed divergence in gene expression in head and larval tissues. Although TEs’ presence does not affect overall gene expression, we observed 6.27% of the expressed genes generating gene-TE chimeric transcripts, including those with function affecting host preference. Our combined genomic and transcriptomic approaches provide evidence of TE-driven divergence between species, highlighting the evolutionary role of TEs in the context of host shift, a key adaptive process that can cause reproductive isolation.

## Introduction

The molecular consequences of host shift evolution can ultimately cause specialization and ecological speciation (1). In many species, reproductive isolation arises as a result of successive adaptations driven by host shift, which results from divergent selection between different environments (1–4). Thus, natural selection increases the frequency of alleles that confer higher fitness benefits when a host shift occurs. *Drosophila* species use a wide range of necrotic host tissues as feeding and breeding sites, providing a suitable model to study the host shift process and its role in insect speciation. Among them, the *repleta* group comprises a clade with ∼100 cactophilic *Drosophila* species (5). The specialization in cacti represents an important ecological challenge to the flies, mainly due to the presence of toxic compounds (6). In the *repleta* species, the host shift was initially to the *Opuntia sp*. cacti, and secondarily, several independent shifts happened to columnar cacti (6), considered a host with higher toxicity than *Opuntia sp*. (7). Such host shift is observed between sibling species with recent divergence in two clusters of the subgroup *mulleri*: cluster *buzzatii* and cluster *mojavensis*. In the cluster *buzzatii, Drosophila buzzatii* and *D. koepferae* are sibling species (divergence 4-5 Mya (8)) with different primary hosts. The former species preferentially use *Opuntia sp*. cacti, whereas the latter is a columnar cacti dweller (9). In the cluster *mojavensis, D. arizonae* and the three allopatric subspecies of *D. mojavensis* have different cacti preferences. *D. arizonae* uses both *Opuntia sp*. and columnar cacti, whereas *D. m. mojavensis* uses the columnar cacti *Ferocactus cylindraceus* as preferential host; *D. m. wrigleyi* uses only *Opuntia sp*., *D. m. baja* uses the columnar cacti *Stenocereus gummosus*, and *D. m. sonorensis* is also a columnar cacti dweller, using *S. thurberi* (10). Therefore, this set of species from *buzzatii* and *mojavensis* clusters provides an excellent model to investigate the molecular mechanisms underlying host shift evolution.

The radiation of cactophilic *Drosophila* species across different hosts is an evolutionary outcome that gave rise to specific adaptations (11). They can be summarized into three steps: localization, acceptance, and host usage (12). In the localization step, odorant-binding proteins (OBPs) and odorant receptors (ORs) are key proteins to distinguish specific hosts in the environment, through the integration of the visual and olfactory systems. Subsequently, in the acceptance step, the insect evaluates the nutritional compounds present on the host, as well as the presence of competitors, predators, pathogens and parasites (12). Several receptors are associated with this process, such as gustatory receptors (GRs) (13) and ionotropic receptors (IRs) (14–16). Finally, the use of metabolites derived from the host is essential for feeding and to complete the breeding process. Although not always associated with detoxification (17), cactophilic flies must overcome the presence of toxic substances from columnar cacti. Several gene families are associated with detoxification, such as Cytochrome P450s (CYPs), Glutathione S-Transferases (GSTs), UDP-Glycosyltransferases (UGTs), esterases (ESTs), and ATP binding-cassette transporters (ABCs). Altogether, these nine gene families associated with the Host Localization, Acceptance, and Usage will henceforth be referred to as HLAU.

Transposable elements (TEs) may play a role in environmental adaptation because of their ability to generate mutations. In most cases, mutations caused by TEs are likely to be deleterious or neutral. Throughout evolutionary time, TEs that remain in the genome tend to be silenced by epigenetic control and/or small RNA pathways (18), accumulating mutations, and losing their transposition ability (19). Despite the majority of TEs becoming silenced, the remaining TE sequences may still contain regulatory motifs or protein domains (20). These sequences can be co-opted by the cell machinery, modifying gene expression or protein sequences of the nearby genes (reviewed in (21)). Depending on the outcome of the cooption for the individual fitness, these TEs can increase their frequency in the population, contributing to populational evolution and hence being determined as adaptive insertions. For instance, in *D. melanogaster*, the gene *CHKov1* has a truncated mRNA derived from a *DOC* TE located into the intron, resulting in resistance to viral infection and insecticides (22). Such exaptation and domestication events can be often identified by the occurrence of chimeric transcripts, which are mRNAs with both gene and TE-derived sequences (23). A recent transcriptome-wide study has identified 327 genes of *D. melanogaster* genes produce chimeric transcripts in different populations (24). Among all genes, 76 generate chimeric transcripts from TE insertions that were present in one strain but absent in another, highlighting the potential of TEs as a source of genetic novelty between different *Drosophila* ecotypes.

Many aspects involving host shift adaptation in *repleta* species have been shown previously, such as detoxification pathways (25), morphology (26), life history traits (27), behavior (28), and genomic and transcriptomic differences (29–33). However, the contribution of TEs to the evolution of HLAU genes has not yet been assessed. Here, we aimed to uncover the extension of the genetic variability derived from TEs in cactophilic *Drosophila* species, using both genome- and transcriptome-wide analysis. We tested the hypothesis whether TEs have contributed to the evolution of HLAU genes, and consequently to the host shift in cactophilic species.

## Results

### Genome assemblies and gene annotation

To investigate the potential role of TEs in host shift of cactophilic *Drosophila* species, we performed Nanopore long-read sequencing on species that have different preferential cacti as hosts: *D. buzzatii* (*Opuntia sp*.), *D. koepferae* (columnar cacti), *D. arizonae* (*Opuntia sp*. or columnar cacti), *D. m. mojavensis* (*F. cylindraceus*), *D. m. wrigleyi* (*Opuntia sp*.), and *D. m. sonorensis* (*S. thurberi*). We obtained high-resolution assemblies for all genomes, with an average of 567 scaffolds, 13.8 Mb of N50 (Supplemental Table 1), and ∼98% of BUSCO genes (Supplemental Fig. S1). Furthermore, the gene annotation for these genomes covered 98.63% of all reference genes in the *D. mojavensis* subspecies, and 95.98% in *D. arizonae*. The *de novo* gene annotation in *D. buzzatii* and *D. koepferae* revealed a total of 18,050 and 17,848 coding genes, for which 63.77%, and 64.13% were successfully assigned as one-to-one orthologs with *D. mojavensis*, respectively. These results represent the higher gene repertoire identified for *D. buzzatii* and *D. koepferae* species, altogether with their ortholog relationship with *D. mojavensis* and *D. arizonae*. Since analysis of HLAU genes is the focus of this study, their efficient annotation is relevant. They are represented by nine gene families associated with the three steps in the use of host resources: (1) Localization: OBPs, ORs; (2) Acceptance: GRs, IRs; and (3) Host usage: ABCs, ESTs, GSTs, UDPs, CYPs (Supplemental Table 2). In *D. arizonae* and *D. mojavensis* subspecies, we recovered ∼98.21% of the total HLAU genes from the reference genome. In *D. buzzatii* and *D. koepferae*, the number of annotated HLAU genes covered ∼95% in both species compared to *D. mojavensis* (Table 1).

**Table 1.**
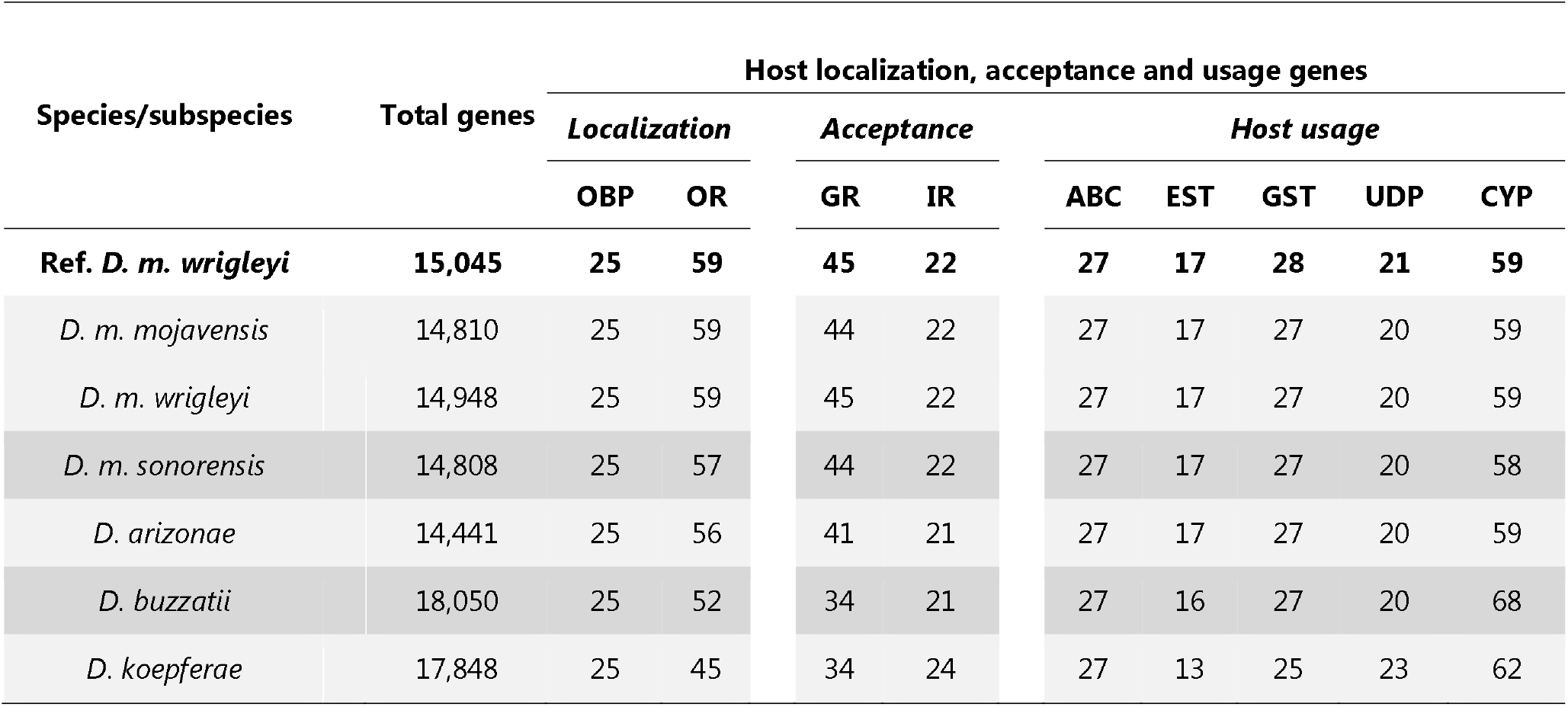
Total number of genes annotated in six *Drosophila* cactophilic species. The “Ref. *D. m. wrigleyi*” represents the annotation in the reference genome used to perform the gene annotation for our long-read Nanopore genomes. HLAU genes are separated according to their functions: Localization: odorant-binding proteins (OBP), odorant receptor (OR); Acceptance: gustatory receptor (GR), ionotropic receptor (IR); Host usage: ATP-binding cassette (ABC), carboxylesterase (EST), glutathione S-Transferase (GST), glycosyltransferase (UDP), and cytochrome P450 (CYP).

### Minor effects of positive selection in HLAU genes

The use of *Opuntia sp*. has been proposed as the ancestral state in cactophilic *Drosophila* species, whereas columnar cacti are the derived state (6). To investigate whether species that prefer columnar cacti as hosts exhibit signs of positive selection in HLAU genes, we employed a branch-site approach to compare their evolutionary rates with those of species that use *Opuntia sp*. For this analysis, we removed *D. arizonae* due to its non-preferential usage between columnar and *Opuntia sp*. cacti (6). Moreover, we added *D. navojoa*, which is a basal species of the *mojavensis* cluster that uses solely *Opuntia* sp. (34). Therefore, we carried out the branch-site selection test with *D. buzzatii, D. m. wrigleyi*, and *D. navojoa* as *Opuntia*-related branches (background), and *D. m. mojavensis, D. sonorensis*, and *D. koepferae* as columnar-related branches (foreground). It is important to highlight that due to the requirement for one-to-one orthologous genes between the six genomes, our analysis has reduced the total number of HLAU genes to 42.9% of the initial number presented in Table 1. From 130 genes, we observed six genes under positive selection (FDR < 0.05): *Obp56d, Obp19d, Gr5a, Ir25a, Cyp4s3*, and *esterase S*. This result suggests that positive selection in HLAU genes is not a major evolutionary driving force in the species under study.

### Evolutionary dynamics of TEs in cactophilic Drosophila species

We used a combination of automatic and manual methods to build a curated TE library for each species, based on the classification of TEs (35). The method was designed to perform polishing steps in both TE consensus and copies to remove artifacts (see Methods).

The total number of TE consensuses (LTRs separated from internal sequences) ranged from 255 in *D. buzzatii* to 372 in *D. m. mojavensis*. The TE content in the genomes ranged from 15.04% in *D. buzzatii* to 22.01% in *D. m. mojavensis* (Fig. 1A). In addition, we observed a positive correlation between TE content and genome size estimated from the assemblies (*Pearson* correlation: *p* = 0.0082; *R* = 0.93) (Supplemental Fig. S2). Both species from the *buzzatii* cluster had the lower genome sizes and TE contents compared to species of the *mojavensis* cluster (Fig. 1A). Regarding the proportional load of TE orders in the genomes, LTRs and Helitrons contribute on average with ∼70% of total TE content in all species (Fig. 1B).

**Figure 1.**
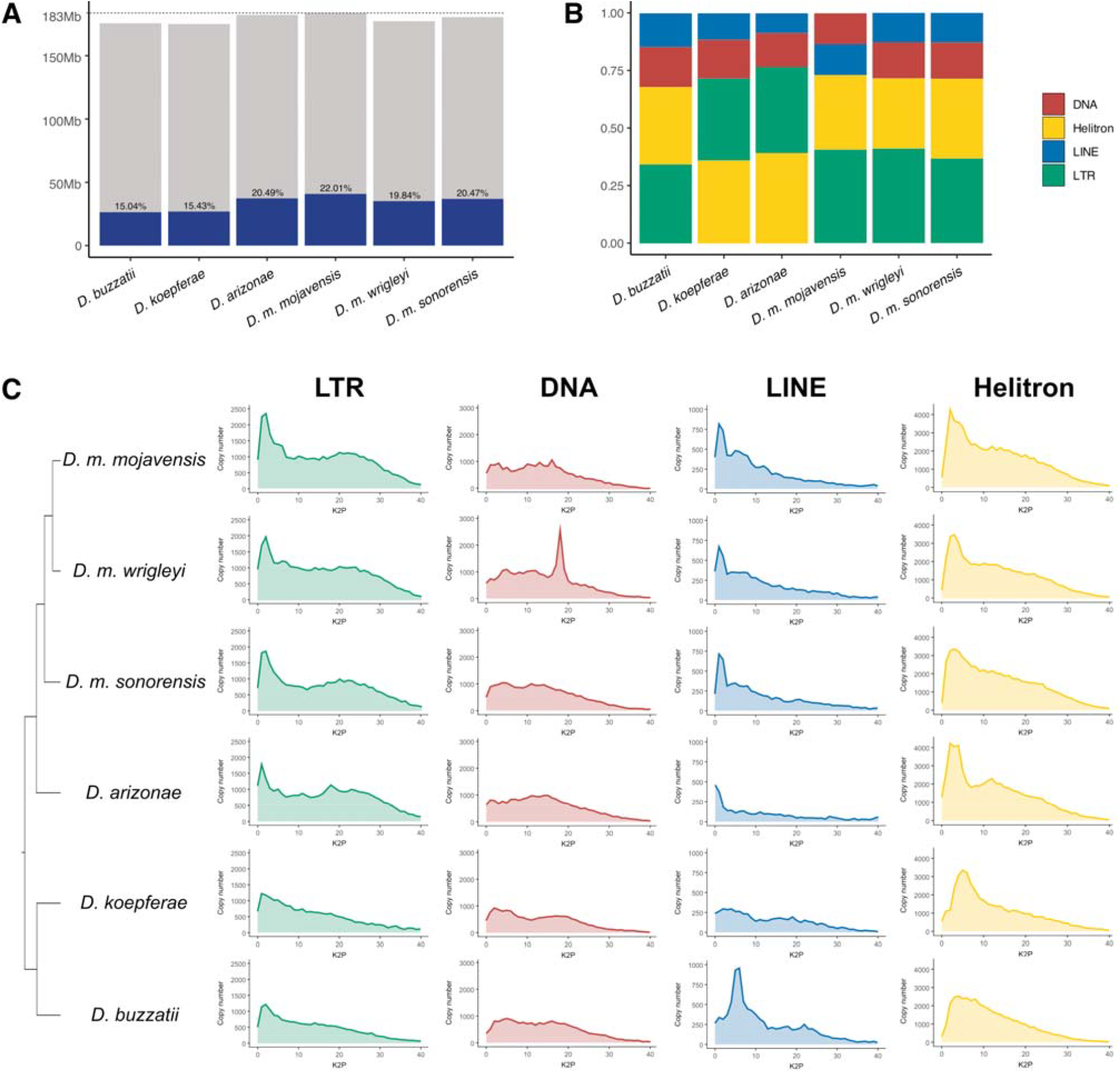
**A**) Genome size of the six cactophilic *Drosophila* species sequenced in this work, and their respective TE content. **B**) Proportional distribution of the total TE content in the genomes, in crescent order. **C**) Phylogenetic relationship among species (adapted from (25,37)), alongside TE order landscapes representing total copy number (y-axis), and the divergence of TE insertions compared to the consensuses (x-axis), based on Kimura 2-parameters (K2P) corrected with CpG. The copies located on the left of the graph have low divergence with their respective consensus, hence inferring that they might be conserved/recent/active, whereas copies on the right side represent copies with high divergence due to the accumulation of mutations, corresponding to old/inactive copies.

In the *mojavensis* cluster (*D. arizonae* and the three *D. mojavensis* subspecies), the abundance of TEs in the genome varied substantially (Fig. 1B), despite their recent divergence time of 1.5 Mya (36). *D. m. mojavensis* has the TE-richest genome (22.01%), with the abundance following from the highest to the lowest: LTR > Helitron > LINE > DNA (Fig. 1B). The distribution of TEs in the other two *D. mojavensis* subspecies and *D. buzzatii* does not follow such order. LTRs and Helitrons have the highest abundance, followed by DNA and LINEs (Fig. 1B). In *D. arizonae* and *D. koepferae*, Helitrons are the most abundant, following LTR, DNA, and LINEs (Fig. 1B). The observed differences are often associated with the divergence in the evolutionary patterns of the TE orders between the genomes. We used copy number as a proxy to measure TE expansion in the genomes, and divergence Kimura 2-parameters (K2P) as age measurement. In the *mojavensis* cluster, we observed that LTRs and Helitrons had a higher expansion in *D. m. mojavensis* compared to the other genomes (Fig. 1C). The higher copy number in *D. m. mojavensis* is mostly from young TEs (K2P 0-5), which might explain the higher TE content in this species. DNA elements have a constant evolutionary dynamics in the genomes, except in *D. m. wrigleyi*. In this species, we observed a peak in copy number with K2P between 16 and 20, considered old insertions. LINEs had a similar evolutionary landscape in *D. mojavensis* subspecies, with higher copy number from young copies compared to *D. arizonae*. In the *buzzatii* cluster, Helitrons show higher copy number from young insertions in *D. koepferae* than in *D. buzzatii* (Fig. 1C), whereas LINEs show a past transposition burst in *D. buzzatii* compared to *D. koepferae*. The antagonistic patterns of TE dynamics observed in the six genomes suggest that TEs had different successful lineages across these species.

### HLAU genes are enriched in TEs within regulatory regions

TE insertions may be selected in the promoter of genes if they provide a benefit to the individual fitness, through the spreading of histone marks, or *cis-*regulatory elements (38,39). To investigate whether TEs are enriched in the promoters (2 Kb upstream of the TSS) from the nine HLAU families, we performed permutation tests in the six genomes (see Methods). To compare our findings with other non-host shift related genes, we also included zinc-finger and serine/threonine kinases as two non-HLAU gene families. We observed that ORs have enrichment of TEs at the promoters in all genomes from species of the *mojavensis* cluster (p-value < 0.05), (Fig. 2A). For acceptance genes, *D. koepferae* had enrichment of TEs in the promoters of GRs (Fig. 2A). Finally, the enrichment of TEs within promoters of detoxification genes was observed in *D. arizonae* (CYPs), *D. buzzatii* (GSTs), and *D. koepferae* (ABCs). The results depicting the histograms with frequency of genes with TEs for each significant permutation test, and p-values are shown in the Supplemental Fig. S3 and Supplemental Table 3. Overall, detoxification genes had z-scores towards depletion of TEs on promoters, with significant depletion for UGTs in *D. buzzatii* (p-value = 0.033) and *D. arizonae* (p-value = 0.0135), GSTs in *D. koepferae* (p-value = 0.0205) (Fig. 2A). In the non-HLAU gene families, we observed the expected result by chance: either z-score indicated the expected number of genes with TEs on promoters in a genome-wide comparison (z-score near 0), or it indicated depletion of TEs, as observed for SER genes in *D. buzzatii*. This result reinforces our findings, indicating the significant and evolutionary unexpected enrichment of TEs in HLAU genes, especially on the promoters of ORs.

**Figure 2.**
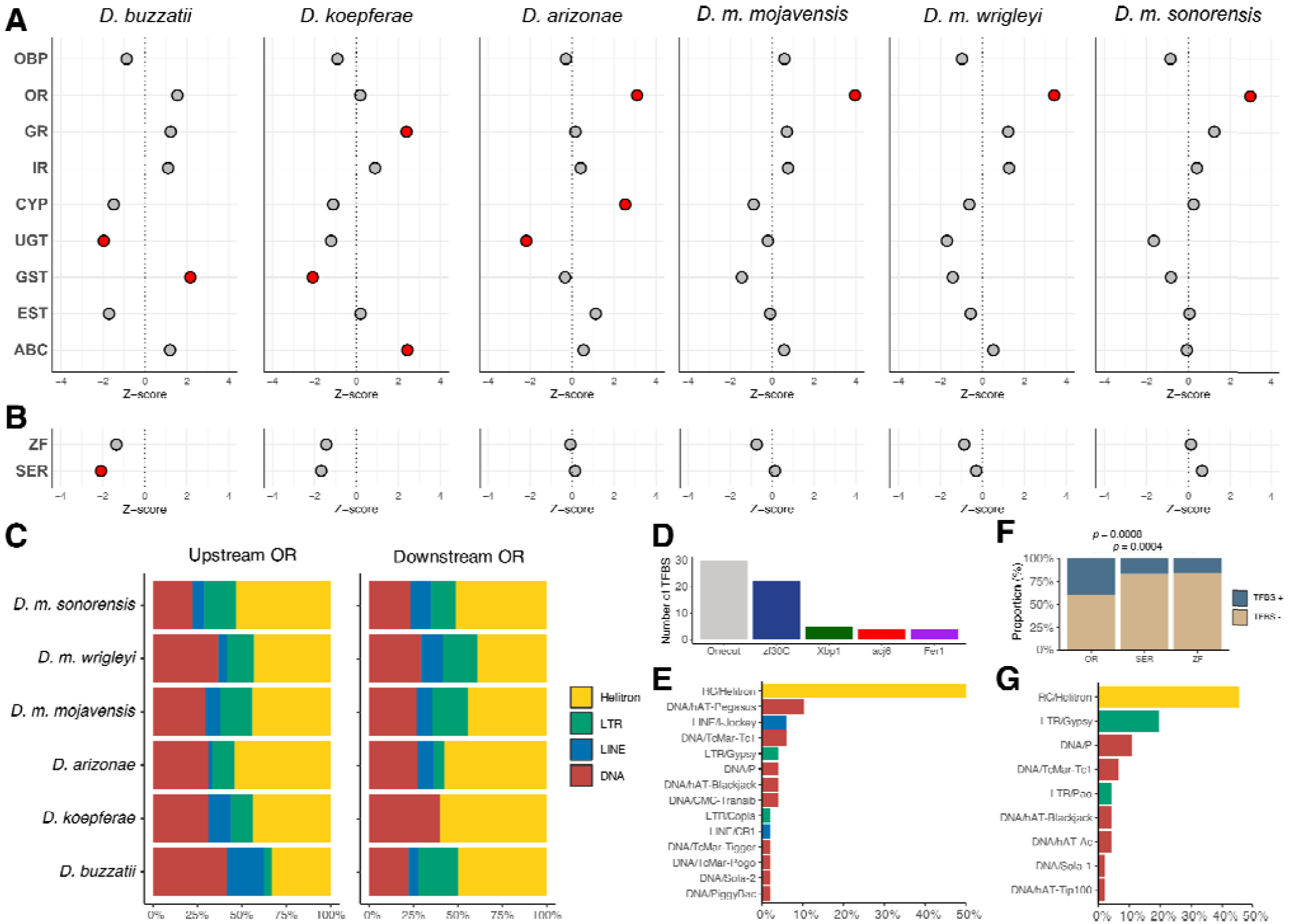
**A**) Enrichment of TEs in the promoter region (2kb upstream) of HLAU gene families. Z-score > 0 indicates enrichment, and Z-score < 0 indicates depletion. Red dots: p-value < 0.05, grey dots: p-value > 0.05. **B**) The same enrichment analysis as depicted on A, but for non-HLAU gene families, represented by zinc-finger proteins (ZF) and Serine/Threonine Kinases (SER). **C**) Proportion of TE orders located upstream and downstream ORs. **D**) Transcription factor binding sites (TFBSs) frequency on TEs located upstream of OR genes in all species under study. **E**) Proportional frequency of TE families with TFBSs embedded on their sequences from TE copies located upstream of OR genes. **F**) Significant Fisher’s exact test between OR genes with OR-related TFBSs from TE insertions compared to the frequency observed in the non-HLAU gene families Serine/Threonine Kinases (SER) and zinc-finger proteins (ZF). **G**) Proportion of TE orders carrying OR-related TFBSs in non-HLAU gene families.

To obtain further insights into the potential regulatory role of TEs, we also analyzed the enrichment at the 2 kb downstream region. Among all species, the only significant enrichments were observed in ORs from species of the *mojavensis* cluster (Supplemental Table 3), as we observed in the promoters. The TE enrichment upstream and downstream of ORs indicates a possible coevolution of this gene family with TEs. Alternatively, it may also suggest that ORs are located in TE-rich regions of the genomes. To investigate this hypothesis, we identified TE-rich regions (Supplemental Fig. S4) and analyzed the frequency of ORs. We observed a single OR in TE-rich regions in *D. arizonae, D. m. mojavensis*, and *D. m. wrigleyi*. (Supplemental Table 4). Therefore, we rejected the hypothesis that ORs are enriched with TEs in their vicinity due to their location in TE-rich regions. Other factors might be involved in the evolutionary maintenance of TEs near ORs, such as the co-option of regulatory motifs.

A few TE families are more prone to contribute to *cis-*elements than others, such as retroelements in *D. melanogaster* (40). Therefore, we investigated whether the observed enrichment of TEs in the HLAU gene families is TE-specific. Considering all species, 45.36% and 50.36% of the insertions located upstream and downstream of ORs correspond to *Helitrons*, respectively (Fig. 2C). We decided to analyze whether these *Helitron* insertions upstream to HLAU genes, mainly ORs, could be part of the first exons from genes, since such overlap has been observed previously in other species than *Drosophila* (41,42). By intersecting exons from genes enriched with their respective upstream *Helitrons*, we observed that *Or85c* (overlap 772 nt) and *Or22c* (52 nt) had overlap with *Helitrons* in *D. m. mojavensis*, whereas *D. m. sonorensis* had only *Or22c* (58 nt). While the insertion in *Or85c* suggests a subspecies-specific insertion, the insertion in *Or22c* is likely to be inherited from the species’ common ancestor (Supplemental Table 5). At the ORs’ downstream region, we observed four genes with *Helitrons* overlapping exons: *Or71a, Or10a, Or13a*, and *Or82a*. Interestingly, the same *Helitron* in the *Or82a* was observed in the three *D. mojavensis* subspecies, highlighting the inheritance from the common ancestor and maintenance of the insertion over time. Taken together, our results rejected the hypothesis regarding the possible role of *Helitrons* carrying HLAU gene fragments. However, these elements could still act on gene regulation if they carry *cis*-elements as TFBSs.

TEs located near genes can modulate gene expression by donating TFBSs or polyA signals to the ancestral regulatory motifs. To investigate the potential functional role of the observed enrichment of TEs in ORs, we performed TFBS identification on all TE insertions located in OR promoters. We used the transcription factors (TFs) previously described to participate in OR’s transcription in *D. melanogaster*: acj6, onecut, xbp1, fer1, and zf30C (43). Taking all species together, the most frequent TFBS found was for the TF onecut (46.15%), followed by zf30C (33.84%), and Xbp1 (7.69%) (Fig. 2D). *Helitrons* were the TEs with the highest frequency, representing 50% of all insertions carrying ORs’ TFBSs (Fig. 2E). Our analysis revealed that 39.28% of the ORs with TEs in promoters have the TFBS located within the insertion (Supplemental Table 6). To test whether these results are specific to ORs, we also performed the same detection of TFBSs and calculation of TEs frequency in the non-HLAU families ZF and SER. Our results show that ZF and SER have, on average, 15.55% and 17.14% of the genes with OR-related TFBSs, respectively. The detection of these TFBSs in ZF and SER is likely due to chance, given the short length of the motifs. Comparing these proportions with the ones found in ORs, we observed significantly more TFBSs compared to ZF (*Fisher’s exact test: p-value = 0.0004341*) and SER (*Fisher’s exact test: p-value = 0.0008519*) (Fig. 2F and Supplemental Table 7). In addition, *Helitrons* also had higher prevalence than other TEs upstream of SER and ZF genes (Fig. 2G), demonstrating a potential insertion preference for promoter regions. Although similar *Helitron* frequencies between OR, SER, and ZF genes, our data suggest that *Helitrons* might have been retained in OR promoters due to the presence of functional TFBSs (Fig. 2F).

### HLAU genes are differentially expressed between Opuntia sp. and columnar cacti dwellers

Differential expression of genes has been associated with adaptation to new hosts in insects (32). Here, aiming to test whether gene expression divergence is associated with adaptation to different cactus hosts, we performed differential expression analysis between species with *Opuntia sp*. preference vs columnar cacti preference, in head and larval tissues. Since our fly stocks were reared in the lab with corn media, the observed transcriptome response is constitutive rather than triggered by the native cacti. Overall, the principal component analysis from HLAU gene expression (normalized read counts) revealed remarkable differences between species in both tissues (Fig. 3A). The variance of gene expression resembles the phylogenetic divergence of the species, with the highest variance between the *buzzatii* and *mojavensis* clusters (PC1), followed by divergence between *D. buzzatii* and *D. koepferae* (PC2). The species of the mojavensis cluster were clustered together with a slight separation between *D. arizonae* and the subspecies of *D. mojavensis*.

**Figure 3:**
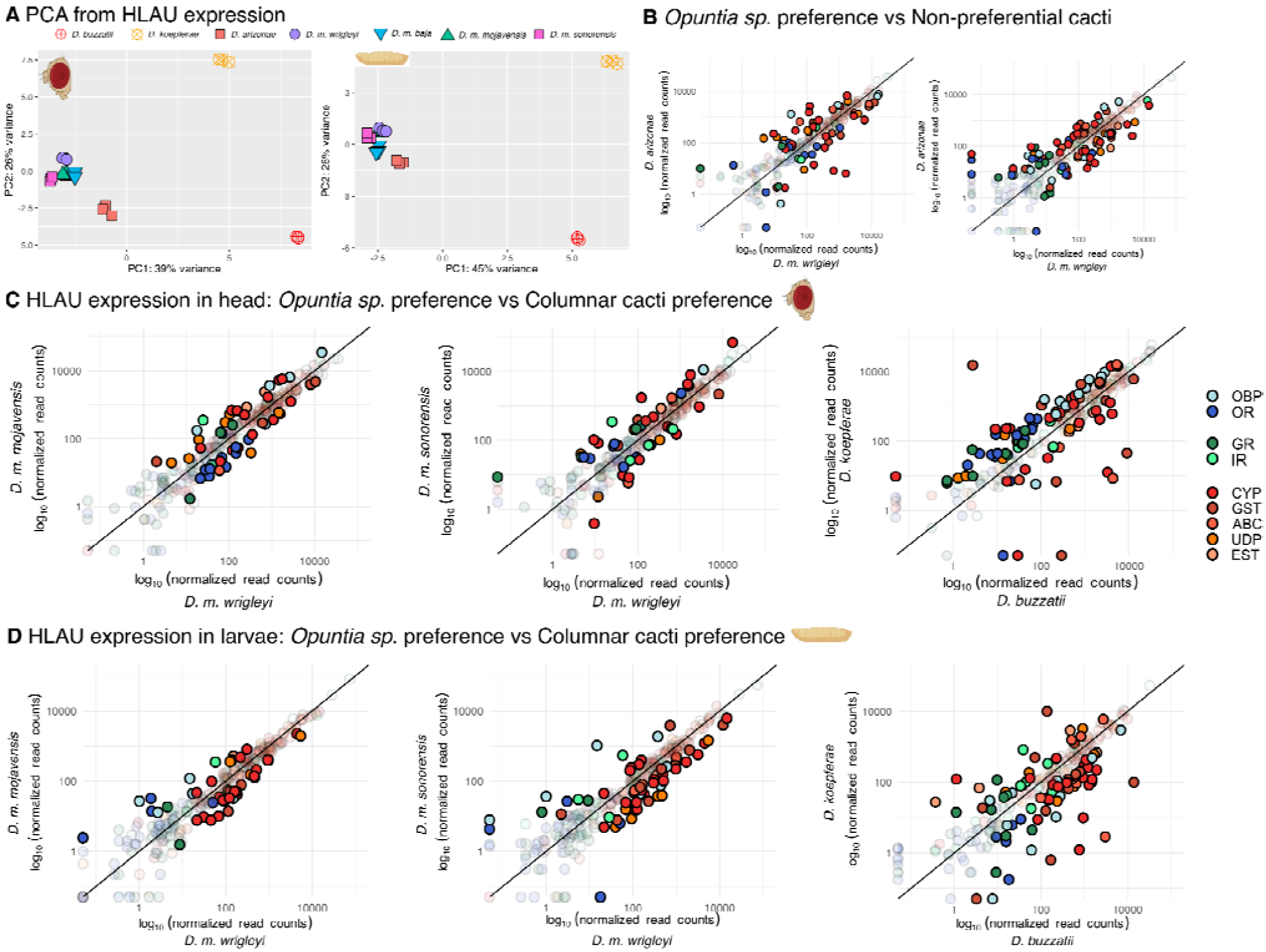
**A**) Principal component analysis of HLAU genes’ expression in head and larval tissues. Both tissues had HLAU genes’ variance representing the phylogenetic distance between species: *D. buzzatii* and *D. koepferae*, compared to the *mojavensis* cluster corresponds to the PC1 (45%), and 26% between them in PC2. Species from the *mojavensis* cluster had strong similarity, with *D. arizonae* slightly separated from *D. mojavensis*. **B**) HLAU genes’ expression in the *Opuntia sp*. dweller *D. m. wrigleyi* versus its sister species *D. arizonae*, which does not have preference between *Opuntia sp*. and columnar cacti. **C**) HLAU expression in head: differential expression analysis of HLAU genes from head in the *Opuntia sp*. dweller *D. m. wrigleyi* versus the three columnar dwellers *D. m. mojavensis*, and *D. buzzatii* versus *D. koepferae*. In A and B: Filled dots represent HLAU genes with log2 fold-change > |1| and FDR < 0.05; transparent dots are non-significant differential expression. **D**) HLAU expression in larvae: The same pairwise species as in A with *Opuntia sp*. dweller vs columnar dwellers, but from larvae tissue.

We performed pairwise analysis of HLAU gene expression between species with *Opuntia sp*. preference (*D. m. wrigleyi* and *D. buzzatii*) versus species with preference for columnar cacti, and *D. arizonae* as non-preferential cacti. All differentially expressed HLAU genes, as well as the presence of TEs in their promoters, can be accessed in (Supplemental Table 8). In head, *D. m. wrigleyi* had 27 and 19 HLAU genes with higher expression compared to *D. m. mojavensis* and *D. m. sonorensis*, respectively (Fig. 3A). Four genes were found with higher expression in the head of *D. m. wrigleyi* compared to the other two subspecies: *UDP-Ugt4, Gst-theta3, Abc-a3*, and *Cyp6a21*. In the subspecies with columnar cacti preference, we observed, on average, 26 HLAU genes with higher expression compared to *D. m. wrigleyi*. Among them, seven were recurrent in the two subspecies: *Ir21a, EstB1, UDP2, UDP-Ugt5, Cyp9b2, UDP-Ugt5-1*, and *Cyp12a2*. Since they were observed in the two subspecies with columnar cacti preference, we propose that they are candidates to be related to the adaptation to columnar cacti. Finally, the same analysis on larvae revealed that *D. m. wrigleyi* has three genes with recurrent higher expression compared to the other subspecies: *Or82a, Obp59a*, and *Ir21a*; whereas in the other two *D. mojavensis* subspecies we observed 17 common up-regulated genes compared to *D. m. wrigleyi*, which comprise seven CYP genes, three GSTs, four UDPs, one Esterase and one OBP (Fig. 3B). Regarding *D. buzzatii* and *D. koepferae*, we observed 27 HLAU genes in head and 54 in larvae with higher expression in *D. buzzatii*, and 63 in head and 29 with higher expression in *D. koepferae*. We also compared the head expression of HLAU genes between *D. m. wrigleyi* and its sister species without cacti host preference - *D. arizonae*. In *D. m. wrigleyi*, we found 31 HLAU genes with higher expression compared to its counterpart *D. arizonae*. In larvae, *D. m. wrigleyi* had 43 HLAU genes with higher expression compared to *D. arizonae* (Fig. 3D). This result demonstrates that HLAU genes have divergent differential expression in both head and larval tissues.

To further investigate whether differential expression of HLAU genes has a higher prevalence compared to non-HLAU genes, we compared the proportion of differentially expressed genes associated with localization, acceptance, and usage with the proportion of differentially expressed genes from the non-HLAU gene families Ser/Thr Kinases and Zinc-finger. In head, our analysis revealed that localization and usage have higher differential expression than non-HLAU genes between *D. buzzatii* and *D. koepferae* (Supplemental Fig. S5). In *D. mojavensis* subspecies, we observed higher differential expression for localization genes between *D. m. wrigleyi* and *D. m. mojavensis*. In larvae, we did not observe any significant difference between non-HLAU and HLAU genes, but the pairwise differential expression between *D. m. wrigleyi* and two subspecies revealed a higher prevalence of gene families associated with usage (detoxification) compared to non-HLAU (Supplemental Fig. S5). In *D. m. wrigleyi*, we also observed higher differential expression of localization genes than in non-HLAU. Finally, the differential expression between *D. m. wrigleyi* and *D. arizonae* demonstrated significantly higher proportion of genes associated with localization and usage compared to non-HLAU in larvae, and higher prevalence of differentially expressed usage genes in head (Supplemental Fig. S5). Taken together, the quantitative results suggest that adaptations that have allowed the shift from *Opuntia sp* to columnar cacti may arise from the evolution of gene expression.

### The enrichment of TEs on HLAU genes is not associated with changes in gene expression

Since we observed enrichment of TEs on the regulatory regions of a few host shift-related families, mainly ORs, and we also found differentially expressed HLAU genes between species, we tested whether the presence of TE insertions is associated with differential expression. To do so, we selected expressed HLAU genes (see Methods) and performed *X*^2^ independence test, splitting them into two variables with two categories each: 1) expression level: differentially expressed, and non-differentially; 2) TE presence: with TEs at 2kb upstream, and without TE at 2kb upstream. Considering all pairwise analysis (Fig. 3A,B-D), the *X*^2^ results demonstrated that the differential expression of HLAU genes is not associated with the presence of TEs on the promoter region (Supplemental Table 9).

Since we did not observe overall effects of TEs on the expression of HLAU genes together, we investigated the association of TEs with differential expression only for HLAU gene families that we found TE enrichment (Fig. 2A). None of the enriched gene families had a significant association between the presence of TEs and their expression level (Supplemental Table 9). Thus, a global enrichment of TEs in the regulatory region is not likely to be associated with the differential expression of HLAU genes.

### Make or break: gene-TE chimeras contribute either predominantly or negligibly to gene expression

Although highly deleterious, a small fraction of TEs generating chimeric transcripts can be positively selected, giving rise to domestication and exaptation events (44). Aiming to identify the extent of the transcriptome variability generated by gene-TE chimeric transcripts, we analyzed the transcriptome of head and larval tissues in the six cactophilic *Drosophila* species. The method employed in this work considered paired-end reads spanning from TEs to exons. Although systematically found in all replicates, many chimeras are likely to be products of pervasive transcription, generating ectopic transcripts whose expression levels are insufficient to play a functional role. Therefore, we developed an automatic method to assemble chimeric transcript isoforms and compute their expression contribution relative to the total gene expression. This method is implemented in the pipeline *ChimeraTE v2.0* (24) as an optional downstream analysis to the total chimeras detection.

Our results revealed that, on average, 715 and 574 genes produce chimeric transcripts across all species, corresponding to 7.51% and 5.04% of the expressed genes in head and larvae, respectively. In both tissues, TE-exonized transcripts represented the highest frequency, with 87.59% in head and 89.09% in larvae (Fig. 4A), followed by TE-terminated transcripts with 10.63% and 9.45%, and TE-initiated transcripts with 1.79% and 1.51%. Interestingly, all species have a similar number of expressed genes(head :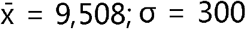;larvae: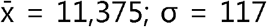), but we observed on average 11-fold fewer chimeric transcripts in *D. buzzatii* and *D. koepferae* compared to *D. arizonae* and *D. mojavensis* subspecies. This result suggests that species from the *mojavensis* cluster are more prone to generate chimeras, or that *D. buzzatii* and *D. koepferae* have a strong purifying selection against gene-TE chimeric transcripts.

**Figure 4:**
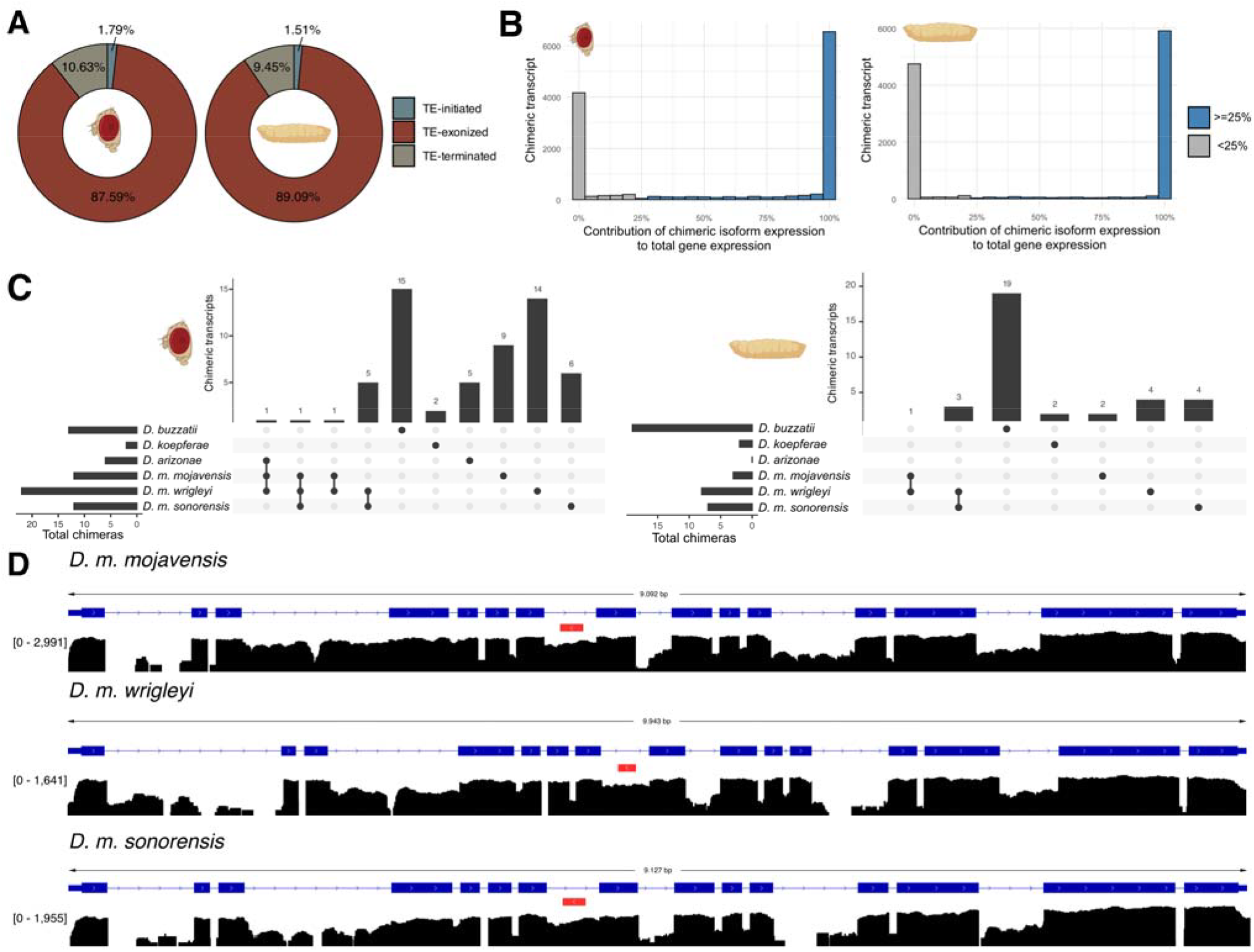
**A**) Proportion of each chimeric transcript category based on the position of TE insertions relative to the gene structure. **B**) Histograms with contribution of chimeric transcript isoform relative to total gene expression (x-axis). Chimeric transcripts without assembled sequence or contribution < 25% were removed (grey bars). **C**) Number of common and unique chimeric transcripts from HLAU genes detected in head and larvae. **D**) *Eato/LOC6578120* (ABC transporter) gene has a conserved TE-exonized transcript with a LINE/R1 element located in its 7^th^ intron. The depth in exons, including the 7^th^ intron, was assessed with RNA-seq from head tissue.

In head and larvae tissues, ∼49% of the detected chimeric transcript isoforms were not assembled or had contribution to gene expression lower than 25% (Fig. 4B). Among these discarded chimeras, the vast majority was not detected by transcriptome assembly approach (0% contribution) (Fig. 4B), which is likely to be associated with negligible expression. Considering chimeric transcripts with contribution >= 25%, more than 95% have contribution equal to 100% (Fig. 4B). This result indicates that almost all genes producing chimeras are dependent on the TE sequence to constitute their complete mRNA sequence. Since ∼88% of all chimeras detected with > 25% contribution are from TE-exonized transcripts, for which 72.35% are from TEs embedded in exons, a high frequency of chimeras with total gene expression could be expected. Notably, this pattern is also observed in TE-initiated and TE-terminated transcripts (Supplemental Fig. S6), despite the TEs being located upstream and downstream of the gene body. Our results reveal a clear binomial trend: chimeric transcripts either have low expression and are unlikely to be biologically relevant, or they account for the total gene expression, suggesting a functional role.

### HLAU genes generate chimeric transcripts

To investigate cases where TEs produce chimeric transcripts from HLAU genes, and their possible role in the evolution of these genes, we identified all chimeras derived from HLAU genes in head and larvae transcriptomes. We found a total of 28 (head) and 24 (larvae) HLAU genes producing chimeric transcripts. None of them are the six genes under positive selection. Notably, ABCs, GSTs, and IRs were the most frequent gene families with chimeric transcripts (Supplemental Table 10). Although *D. buzzatii* had 11-fold fewer chimeras than species from the *mojavensis* cluster, we observed 29 gene-TE (a few genes with more than one chimera) chimeras derived from HLAU genes in both tissues, the highest number among all species. Its sister species *D. koepferae* had only four chimeras, which are the same two ABC genes in head and larvae (Supplemental Table 10). In *D. buzzatii*, ∼93% of all chimeras are from GSTs, expressed in both larvae and head tissue. Interestingly, all of them represent cases of *Helitron* exonizations (Supplemental Table 10). We did not find any GST with chimeric transcripts in the other transcriptomes, revealing an unprecedented recent evolutionary novelty in GSTs’ evolution in *D. buzzatii*.

Chimeric transcripts from HLAU genes are represented by 14.81% TE-terminated transcripts and 85.19% TE-exonized transcripts. We did not observe any TE-initiated transcript from HLAU genes. Despite the compelling evidence of TEs enrichment in the promoter of OR genes from species of the *mojavensis* cluster, we did not observe chimeric transcripts from ORs at high frequency. Only *Or45b* (*LOC6578440*), *Or13a* (*LOC26527676*), and *Or85a* (*LOC26528431*) had chimeric transcripts in *D. m. sonorensis, D. m. wrigleyi*, and *D. arizonae*, respectively (Supplemental Table 10). This result reveals that TEs located in the promoter of ORs are not likely to be co-opted as alternative transcription start sites. In the species of the *mojavensis* cluster, we observed ABCs as the gene family with the highest frequency of chimeras (Supplemental Table 10), despite not having enrichment (Fig. 2A). We observed, on average, 5 ABC genes with exonization of multiple TE copies in this group of species. This result indicates that the genomic distribution of TEs is not necessarily associated with their transcriptional interaction with genes.

The quantitative analysis on chimeric transcripts revealed transcriptional polymorphism in HLAU genes due to TEs. We investigated common chimeras present in more than one species, highlighting their retention over evolutionary time, as well as species-specific chimeras that may be associated with recent insertions. Our results revealed that 94.44% and 88.57% of the chimeric transcripts derived from HLAU genes are species-specific in head and larvae tissues, respectively (Fig. 4C). In head, the *Opuntia sp*. dwellers *D. buzzatii* and *D. m. wrigleyi* were the ones with the highest number of HLAU genes with species-specific chimeric transcripts (Fig. 4C). The 15 chimeras uniquely found in *D. buzzatii* are derived from 12 genes (2 ABCs and 10 GSTs); whereas the 14 chimeras from *D. m. wrigleyi* were observed from 10 genes (5 ABCs, 3 IRs, 1 CYP, and 1 OBP) (Supplemental Table 10). In larvae, *D. buzzatii* follows a similar result as observed in head, 15 chimeras derived from 14 genes (2 ABCs and 12 GSTs), while *D. m. wrigleyi* had only four unique chimeras (2 ORs, 1 IR, and 1 CYP).

Cases of common chimeric transcripts between populations/species may indicate exaptation/domestication events. In the six transcriptomes analyzed here, we found 8 genes in the head and four in larvae present in more than one genome (Supplemental Table 11). These common chimeric transcripts occurred only between *D. mojavensis* subspecies, revealing that TEs interacting with HLAU genes are likely to be lost over higher evolutionary scales. Their inclusion into the mRNA sequence and high contribution can be observed. For instance, the ABC transporter (*Eato* gene) has an exonized LINE/R1, which is located in the 7^th^ intron (Fig. 4D), with an average contribution of 84.25% to the gene expression. Taken together, these results demonstrate unprecedented transcriptional interaction between TEs and HLAU genes in a tissue-specific manner.

## Discussion

Insect host shift and its role in speciation have been the subject of research to understand the evolution of reproductive isolating barriers and adaptation to local environments. Indeed, many insects have finely tuned adaptations to locate, feed, reproduce, and develop on the host tissue. Adaptations to new hosts can lead to reproductive isolation, wherein ecological speciation arises as a consequence within divergent natural populations inhabiting distinct environments. In this context, we investigated whether TEs could play a role in contributing to the evolution of HLAU genes, with the potential to facilitate rapid adaptive radiation to new hosts. To test this hypothesis, we selected six *Drosophila* species/subspecies from the *repleta* group with different primary cacti hosts.

### Weak signal of positive selection in HLAU genes through branch-site model

The necrotic tissue of each cactus-host used by cactophilic *Drosophila* species differs in its composition in terms of yeasts and alkaloids (45,46). Therefore, species have adapted to feed and breed on different hosts, as demonstrated by behavioral and physiological response when species feed and breed in non-preferential hosts. Positive selection in HLAU genes have been proposed to allow the recognition of a broader spectrum of odors, metabolize nutrients, and activate detoxification pathways (12). Indeed, specialist species are likely to lose some of their genes related to sensory pathways, while the genes that are retained are subject to positive selection (47,48). A preliminary genome-wide analysis of signatures of positive selection in the four *D. mojavensis* subspecies revealed high ω rates on genes associated with chemosensation, perception, immunity, behavior, detoxification, and reproduction (49). Although a previous genome-wide analysis has identified 1,294 genes under positive selection (50), here we focused the analysis to investigate specifically HLAU gene families. Our results with brach-site model had only six HLAU genes under positive selection: *Obp19d, Obp56d, Gr5a, Ir25a, Cyp4s3, esterase S*. The unprecedent method used in this work focused on the common differences of species adapted to columnar cacti, rather than independent evolutionary events, making it more stringent. The lack of genes with statistical support for positive selection may be also associated with technical limitations, since we based our analysis on 1:1 orthologs, and a certain proportion of the genes evolve too fast to be correctly assigned during orthologs identification (51). Additionally, we observed ω values higher than 1.5, for instance as *Gr59d* with ω = 3.49. Although we did not observe signficiant positive selection, this result indicates ongoing divergence between species/subspecies.

### OR promoters are enriched with TE insertions

The literature has recurrently demonstrated that TEs can create new regulatory elements, including gene promoters (52–54), enhancers (55,56), and insulators (57,58). In mammals, TE-derived sequences contribute up to 40% of genome-wide binding sites for transcription factors located in TE-rich regions (59). It has been proposed that such co-option of TE-regulatory elements for gene regulatory networks can provide substantial modification of gene expression over short time scales, contributing to genetic variability at the transcriptome level (60–62). The colonization of new habitats and hosts is accompanied by behavioral changes, mainly through alterations in environmental perception (63,64). Here, we observed that ORs are enriched by TEs in the four species from the *mojavensis* cluster. In ants, ORs have also been found enriched in TEs. But in ants, TEs are associated with tandem duplication by crossing over, allowing massive duplication events (65). Since there is no expansion of ORs in cactophilic species, the enrichment in TEs is likely to have another function, probably related to gene regulation due to its specific location in promoters. Although we did not observe a significant enrichment of TEs in differentially expressed genes, we argue that TEs might contribute to the regulation of specific HLAU genes. In ORs, ∼39% of the genes have OR’s TFBSs located within TEs. We propose that this subset of genes is not sufficient to have a significant signal when tested with ORs without TEs, but they may have functional TE-derived *cis*-regulatory elements.

*Helitrons* have been extensively reported on the promoter of other eukaryotic species (41,66–68), in which a few cases of embedded TFBSs were reported (69,70). Here, we demonstrated that ∼50% of the enrichment in TEs observed at the promoter region of ORs might be due to the exaptation of TFBSs derived from *Helitrons*. In addition, the recurrence of the enrichment in species of the *mojavensis* cluster might demonstrate potential cases of TE co-option inherited by the common ancestor. Our genomic analysis provides a robust list of potential candidate genes in which *Helitrons* might play a regulatory role. This is an interesting hypothesis that can be determined through cutting-edge technologies such as CRISPR methods to delete insertions with site-specific resolution (71). Our results suggest a putative role of TEs in the regulation of HLAU genes across multiple species/subspecies.

### Head transcriptome reveals subspecies-specific patterns

The differential expression of HLAU genes in the head tissue revealed that localization and host usage (detoxification) gene families have the highest divergence across the subspecies, since we observed more differentially expressed genes for these families. In *D. m. wrigleyi*, which is the only *D. mojavensis* subspecies with a preference for *Opuntia sp*., we observed that CYP genes had on average 12 up-regulated genes compared to other species. This result highlights that the divergence of the cacti preference between *D. mojavensis* subspecies might be explained by modifications in gene expression of HLAU genes in the head tissue. Although *Opuntia sp*. has lower toxicity than columnar cacti (72), we found a similar number of up-regulated detoxification genes between *D. m. wrigleyi* and the other subspecies. This suggests that *D. m. wrigleyi* has a constitutive detoxification response that is independent of host toxicity. This is a plausible explanation since *Opuntia sp*. produces fewer benzaldehyde volatiles than the columnar cacti *S. thurberi*, but both cacti are lethal to *D. melanogaster* after 48 hours (72). Therefore, the adaptation to use *Opuntia sp*. must have a constitutive gene expression of detoxification pathways in *D. m. wrigleyi*.

Comparative transcriptomic studies have revealed divergence in gene expression between *D. buzzatii* and *D. koepferae*, particularly in head tissues associated with sensory perception and detoxification. For instance, Guillén et al. (2014) reported significant differences in the expression of genes involved in olfaction and xenobiotic metabolism, suggesting that these species have adapted to their respective cactus hosts (*Opuntia* for *D. buzzatii* and columnar cacti for *D. koepferae*). Similarly, divergence in expression patterns has been reported in these species in ORs, OBPs, and cytochrome P450 genes (73). These reported differences are consistent with our results in head and larval tissue, reinforcing the hypothesis that ecological specialization and divergence in head transcriptomes play a role in host shift between *D. buzzatii* and *D. koepferae*.

Previous studies with *D. melanogaster* have shown that sequence divergence in ORs and OBPs significantly modulates odor sensitivity (64,74), potentially leading to divergence of host location. In addition, many ORs are involved in a broad range of ecological interactions, influencing behaviors associated with oviposition and feeding (75). For instance, OR evolution has mediated the herbivorous nature of leaf-mining specialist *Scaptomyza flava* (76). Furthermore, transcriptomic analyses have identified 18 ORs that are differentially expressed between heads of *D. m. wrigleyi* and *D. m. mojavensis*, demonstrating that modifications in the sensory transcriptome have evolved between these species (77). In this study, we expanded the understanding of this divergence by comparing species with a preference for *Opuntia sp*. and columnar cacti, and showed that each species has a distinct set of differentially expressed HLAU genes.

### Gene expression in larvae is one of the factors shaping the adaptation to new hosts

Differences in larval behavior and survival have been observed when *Opuntia*-related species are reared in columnar cacti (78,79). A previous study showed that *D. m. mojavensis*, which primarily uses *S. thurberi* as its host, has a 60% reduction in larval viability when reared on *S. gummosus*-based media (80). The authors also proposed that 21% of the genes are differentially expressed between larvae reared in the primary and alternative cacti hosts. Many genes were reported with detoxification, energy production, among others. Another similar study with larvae transcriptome, but with *D. buzzatii* and *D. koepferae*, proposed that *D. buzzatii* flies reared in columnar cacti media have a stronger detoxification response in comparison with *D. koepferae* flies, suggesting that *D. koepferae* has a canalized transcriptome response to the toxic alkaloids from columnar cacti (81).

Here, we observed that the majority of differentially expressed HLAU genes are from gene families associated with detoxification. Since *D. m. wrigleyi* and *D. buzzatii* use primarily *Opuntia sp*., we expected to find a higher number of detoxification genes in the other species/subspecies with columnar cacti preference. However, we observed a similar number of differentially expressed detoxification genes regardless of the host preference. CYPs were overrepresented in our results, but they represent a diverse gene family that can have functions other than detoxification, such as those related to developmental pathways in larvae (82,83). Thus, further analysis of the biological role of up-regulated CYP genes in *D. m. wrigleyi* and *D. buzzatii* might be assessed. Alternatively, as we observed in the head transcriptome, although *Opuntia sp*. has lower toxic compounds compared to columnar cacti, *D. m. wrigleyi* and *D. buzzatii* may also have a resistance to toxicity (72). Thus, any cactophilic species is expected to have evolved a certain level of detoxification ability, which explains our observations. In this study, we propose a set of common HLAU genes in columnar cacti dweller species associated with detoxification. We observed that the three *D. mojavensis* subspecies have the same seven genes with higher expression compared to *D. m. wrigleyi*. Most of them are associated with detoxification (*EstB1, UDP2, UDP-Ugt5, Cyp9b2, UDP-Ugt5-1, Cyp12a2*), except *Ir21a*. It is noteworthy that other evolutionary forces than gene expression might be associated with host shift. For instance, gene duplication events have been reported in CYPs and GSTs in cactophilic species (84). These duplications are thought to enable flies’ resistance to different toxic environments. Similarly, *D. buzzatii* and *D. koepferae* show both copy number variation and expression divergence in detoxification genes, indicating that gene duplication followed by regulatory or coding sequence divergence contributes to host-specific physiological adaptations (50,84). These findings support the view that gene duplication serves as a key evolutionary mechanism facilitating ecological specialization in cactophilic *Drosophila*. Our results show that divergence in the regulation of gene expression in larvae is one of the factors likely to shape the adaptation to new hosts.

### Transposable elements are a source of transcriptome variability between cactophilic species

TEs contribute to genome and transcriptome evolution by rewiring regulatory networks, and by incorporating protein domains into gene transcripts. In both cases, TE copies that are inserted within genes can integrate their sequences into the mRNA, generating chimeric transcripts. A recent study with different ecotypes of *D. melanogaster* has shown that these chimeras have the potential to generate transcriptome novelties (24). These chimeras have also been demonstrated to be active in a tissue-specific manner (85). Although the production of chimeric transcripts derived from polymorphic TE insertions is well-known, there is no study addressing the question of how TEs contribute to the evolution of genes associated with host shift. Here, we took advantage of genomic and transcriptomic data to analyze the transcriptional interaction of TEs with HLAU genes through gene-TE chimeric transcripts. Our finding demonstrated that around half of the chimeras have low expression levels (0-5%), and are likely to represent pervasive transcription of TEs. Notably, over 95% of the chimeras overpassing the 5% contribution to gene expression constitute nearly the total gene expression.

Considering previous findings in different tissues of *D. melanogaster*, where 38% of all chimeric transcripts express only the chimeric isoform (85), our result shows a higher prevalence of chimeras in the repertoire of gene splicing. In tetrapods, the exonization of TE-derived transposases has been shown to constitute the unique gene isoform (86). In HLAU genes, all chimeric transcripts represent TE-exonized transcripts. We suggest that TEs in the promoters of ORs might still recruit transcription factors at their terminal sequences, which would prevent their inclusion in the gene mRNA (chimera). In addition, the maintenance of TEs in the promoters of ORs might also be due to epigenetic marks. TEs can have chromatin marks (87,88), making them versatile regulatory motifs acting in the regulation of nearby genes (89,90). Further experimental analysis, as ChIP-seq, may be performed to test the TEs’ function in OR’s transcription. Furthermore, TE annotation remains challenging in non-model organisms. Although we performed several steps to remove potentially false TE copies from the genome, we can not rule out the possibility of remaining annotation errors in the exons of HLAU genes.

A comparative analysis between generalist and specialist *Drosophila* species revealed that niche amplitude is not likely to play a role in TE dynamics in the genomes (91). Nevertheless, several studies have proposed that the successful adaptation to new environments in invasive species might be associated with TE activity, despite reduced genetic diversity caused by founder effects (92,93). Here, we demonstrate that TEs interact with genes associated with HS, providing a transcriptomic variability in six cactophilic species with different host preferences. Our results show that ABC transporters produce chimeric transcripts in all cactophilic species, highlighting a compelling mechanism of structural innovation. In insects specifically, MITEs (non-autonomous DNA transposons) have been repeatedly identified within defensome genes, including ABC transporters, such as in *Helicoverpa* spp., suggesting a potential causal link to insecticide resistance (94). Functionally, exonized TEs are stably expressed and translated, often producing altered protein domains or localization signals, thereby expanding proteomic repertoires while providing raw material for adaptive selection (95). Taken together, our findings suggest that TE exonization in ABC transporters may offer cactophilic *Drosophila* a versatile genomic strategy to fine-tune transporter function, potentially under the selective pressures of xenobiotics. Although we found a high contribution of the chimeric isoform to total gene expression, future work should be conducted to elucidate which exonized TEs are translated, and whether they modulate substrate specificity, protein stability, or cellular localization, thereby affecting organismal fitness.

Our discovery of 13 GST genes in *D. buzzatii* harboring exonized Helitrons underscores an unprecedented finding of TE-driven functional diversification in detoxification enzymes. *Helitrons* have been shown to supply novel exons, splice sites, and promoters across eukaryotes (41). This finding, unique to *D. buzzatii*, suggests a lineage-specific co-option of *Helitrons* to expand and diversify GST isoforms. GSTs are central to xenobiotic metabolism, and exonization of Helitron-derived sequences could introduce novel domains or alter enzymatic properties, potentially conferring adaptive advantages under environmental pressures such as exposure to cactus-derived toxins. Future work should be performed to determine the function of these *Helitron*-derived motifs in the host preference of *D. buzzatii*.

It is important to state that the identification of chimeric transcripts through transcriptome analysis is an important, but limited step in terms of function. These mRNA molecules might be degraded by surveillance pathways that control for aberrant mRNA production, such as no-go decay (96), non-stop decay (97), and nonsense-mediated RNA decay (98). Our study provides a set of HLAU genes producing chimeric transcripts, in which part of them might have a phenotypic impact and ultimately be associated with host preference. However, it is fundamental to perform comparative phenotypic studies of host adaptation cues with mutant strains for the TEs interacting with HLAU genes identified in this study. The application of techniques of RNAi aiming at chimeric transcripts, or deletion of TE insertions related to chimeras with CRISPR/Cas9 (71) would be crucial to confirm the predictions of our results.

### Transposable elements driving host shift

Host specificity is a key mechanism of reproductive isolation in plant pathogens and is regulated by the repertoire of effector genes within each pathogen. In *Phytophthora sp*., the host specificity is partly regulated by RXLR class effectors that facilitate host exploitation. Notably, synthetic chimeras of a short interspersed element (SINE) linked to an effector gene in *P. infestans* induced their silencing (99). This silencing likely occurs naturally, as transcriptional inactivation of effectors is known to occur, and over half of RXLR effectors in the *P. infestans* genome are located in TE-rich regions (100). Consequently, TEs inserted near these genes may have influenced host shift in *P. infestans*.

Our findings demonstrate multiple interactions between genes and TEs, highlighting the contribution of TEs to the genome and transcriptome evolution of cactophilic species. Although our data did not allow the quantification of the TE insertions frequency in natural populations, our results demonstrate the first transcriptome-wide evidence of TE co-option in cactophilic species. The genes producing chimeras with TEs must be further studied, since the lack of gene-phenotype association between HLAU genes is not fully reported, preventing us from pinpointing a specific relation between chimeras and possibly host preference. In addition, our study focused exclusively on nine gene families that have been shown to evolve during host shift events (12). However, host shift is undoubtedly a complex trait to study, and many other genes can be associated with the adaptation to new hosts. For instance, differential expression of genes associated with development and neurological processes has been identified in cactophilic species reared in different cacti media (101). Behavioral changes are also based on host shift evolution and have been reported in many systems (102). Despite focusing on nine relevant gene families, we suggest further studies including other aspects of host shift than localization, acceptance, and host usage.

The existence of reproductive barriers between populations that undergone a host shift is the key process to ecological speciation. In our model, with species from the *buzzatii* and *mojavensis* clusters, incipient reproductive isolation has been reported between *D. m. sonorensis* and *D. m. baja* (26,103,104). The premating isolation between them occurs due to the difference in cuticular hydrocarbons, which participate in the signaling pathway of mate recognition (105). Importantly, artificially shifting the cactus host was the main cause of changes in the hydrocarbon profiles. Thus, the source of reproductive barriers is dependent on the cacti species on which the flies feed and breed (106,107). The close relationship between HLAU genes to the host shift process, and the observed contribution of TEs to the evolution of these genes, supports a possible role of TEs in the host preference of cactophilic species.

## Methods

### Fly stocks and genome sequencing

Most of the strains were obtained from the UC San Diego *Drosophila* Stock Center: *[D. m. mojavensis*: Anza (01), Anza Borrego Desert, California, USA, Stock Center n°: 15081-1352.01; *D. m. wrigleyi*: CI (22), Catalina Island, California, USA, Stock Center n°: 15081-1352.22; and *D. m. sonorensis*: AG (26), Agiabampo Bay, Sonora, México, Stock Center n°: 15081-1352.26] and *D. arizonae* [HI (17), collected in Metztitlan, Hidalgo, Mexico, Stock Center n°: 15081-1271.17]. D. *buzzatii* (Bu28) and *D. koepferae* (Ko2) stocks correspond to two laboratory lines used in previous works (108,109). Both stocks were maintained by brother-sister mating for more than a decade and then kept by mass culturing.

We used the same protocols of DNA extraction and genome sequencing for all samples. DNA was extracted from 10 males and 10 females from each species using the Qiagen DNeasy Blood&Tissue kit. Then, we evaluated the genomic DNA amount and quality with NanoDropTM One UV-Vis spectrophotometer (Thermo Fisher Scientific, Waltham, MA, USA) and Qubit ® 1.0 Fluorometer (Invitrogen, Carlsbad, CA, USA). Three micrograms of DNA were repaired using the NEBNext FFPE DNA Repair Mix (NEB M6630). We performed end repair and dA-tailing using the NEBNext End repair/dA-tailing Module (E7546, NEB). Ligation was then assessed with the Ligation Sequencing Kit 1D. MinION sequencing was performed according to the manufacturer’s guidelines using R9.4.1 flow cells (FLO-MIN106, ONT) and a Nanopore MinION Mk1b sequencer (ONT) controlled by *ONT MinKNOW v.18.3.1*. Base calling was performed after sequencing using *Guppy v.4.0.5* in high accuracy mode for isogenic wild-type strains *v.3.1.5*.

### Genome assemblies and gene annotation

The quality control for Nanopore reads was performed with *Nanoplot* v.1.10.2 (https://github.com/wdecoster/NanoPlot). Reads with quality lower than 7 were removed from downstream analysis. In order to assemble contigs, *Flye v.2.8* (110) was used with default parameters, except –plasmids. All contigs were aligned with *minimap2* v2.16 (111), with –x map-ont option. The alignment was used to perform the polishing of contigs with four rounds of *RACON* v1.3.2 (112), default parameters. The assembly quality metrics was assessed with *Assembly-Stats* v1.0.1 (https://github.com/sanger-pathogens/assembly-stats). Assembly incongruences were manually visualized with *D-genies* v1.2.0 (113) and corrected with *samtools* v1.9.0. (114), function *faidx*; and *Gepard* v1.4.0(115) for determination of breaking points. The super scaffolding of all corrected assemblies was performed with *RaGOO* v.1.1 (116), with –s and –t 4 parameters, using the respective reference genome assembly for each species. Subsequently, the chromosome-scale assemblies were submitted to a Benchmarking Universal Single-Copy Orhologs (BUSCO) v.5.4.3 analysis (117) with lineage *diptera_odb10* (*-l* parameter) with nucleotide sequences. The gene annotations for *D. mojavensis* subspecies and *D. arizonae* were performed with the software *Liftoff v.1.6.3* (118), with the reference genome of *D. mojavensis* (GCA_018153725.1) (119). We have used the parameters *–exclude_partial*; *-a 0.5* and *–s 0.75* to keep annotated genes with at least 75% of the reference gene length. We removed from our gene annotation HLAU genes flagged as “low quality protein” and “unestablished gene function” in the reference *D. mojavensis* genome.

*D. buzzatii* and *D. koepferae* genomes were annotated with a de novo strategy with the software *BRAKER3 v.3.0.8* (120). To optimize the prediction of gene structures and repertoire of isoforms, we used a mix of RNA-seq data from different tissues publicly available and produced in this work (Supplemental Table 12), in addition to protein sequences from *D. mojavensis* as reference. Ortholog genes between *D. buzzatii, D. koepferae, D. mojavensis*, and *D. arizonae* were obtained with *Orthofinder v.2.5.4* (121) using *DIAMOND v.0.9.14* (122) as search engine and default parameters. One-to-one orthologs were obtained directly from *OrthoFinder v.2.5.4*, and remaining proteins without an ortholog pair were analyzed with *blastp v.2.5.0+* (123), for which we considered as one-to-one proteins only those with a single homologous with parameters *-qcov_hsp_perc 95* and identity >= 90%. Since *Orthofinder* performs its search with protein sequences, the identification of orthologs from non-coding RNA genes was assessed with *Liftoff v.1.6.3* (124) using all non-coding RNA genes from *D. mojavensis* as reference to maximize our homology-based gene annotation.

### Molecular evolution analysis

To identify HLAU genes with signatures of positive selection in species that prefer columnar cacti compared to those that prefer *Opuntia sp*., we selected the three *D. mojavensis* subspecies, *D. navojoa, D. koepferae*, and *D. buzzatii. D. arizonae* species was removed from this analysis due to its generalist preference for both columnar and *Opuntia* cacti. The *D. navojoa* genome was retrieved from NCBI (GCF_001654015.2) (125). For the other Nanopore genomes sequenced in this work, we performed a polishing step with Illumina paired-end reads. We used the software *NextPolish* (126) to fix base errors in the assembled genomes. The three *D. mojavensis* subspecies were polished with genomic Illumina reads sequenced previously (127), whereas for *D. koepferae* and *D. buzzatii*, we used publicly available genomic Illumina reads (128) and RNA-seq reads (50), respectively. In the latter, to increase the read coverage and optimize the polishing to the highest number of genes and exons as possible, we merged RNA-seq reads from adult males and females (50). All genomes were corrected using two rounds of polishing, as recommended (126). Then, we extracted the longest CDS sequences for all genes combining AGAT (129) *agat_extract_sequences.pl* parameters *-t mrna, samtools faidx* (130), and *seqtk* (https://github.com/lh3/seqtk). Posteriorly, we intersected the occurrence of orthologous genes between the six genomes, recovering a list of genes that were present in all of them. The sequences were aligned using *MAFFT v.7.487* [67] and the non-aligned codons were removed using *pal2nal* (131). To obtain a strongly supported tree for this analysis, we inferred the species phylogeny tree with 2,043 shared BUSCO genes. The phylogeny was estimated using the concatenated alignment (3,730,914 sites) with *IQ-TREE v. 2.2.2.6* (132), using the best model for each gene, 10,000 replicates of ultrafast bootstrap and 1000 replicates for Approximate Likelihood-Ratio Test (ALRT) (parameters *-m MFP -bb 10000 - alrt 1000*). Finally, we carried out an analysis to detect signatures of positive selection for each gene individually with *CODEML* from *PAML v.4.10.7* (133), using *branch-site* approach. In order to find signatures of positive selection in species that use columnar cacti, we marked the respective branches of *D. koepferae, D. m. mojavensis*, and *D. m. sonorensis* with the same label (#1), to compare them with the background species that use *Opuntia sp*. (*D. buzzatii, D. m. wrigleyi*, and *D. navojoa*). For this, we tested two hypotheses: (i) H0 – Null hypothesis, in which all branches evolved under negative selection (model = 2, Nssites = 2, fix_omega = 1, omega = 1); (ii) H1 – Alternative hypothesis, in which the labeled branches (foreground) have sites with signatures of positive selection (model = 2, NSsites = 2, fix_omega = 0, omega = 1.5). The hypotheses were tested through a chi-square test, calculated by the log-likelihood (lnL) ratio between H1 and H0 (2ΔlnL) and degrees of freedom obtained from the difference between the number of estimated parameters for H1 and H0 (Δnp). Sites under positive selection were identified by Bayes Empirical Bayes and posterior probability, implemented in *CODEML*.

### RNA extraction and sequencing

We performed RNA-seq for larvae and head tissues from *D. buzzatii, D. koepferae, D. arizonae*, and the four *D. mojavensis* subspecies. The RNA extraction was assessed with the RNeasy Plus Kit for 10 female heads (10 days old) and 15 larvae, with three technical replicates. The library preparation was done with stranded mRNA-seq/standard quantity (ligation), and 100 bp paired-end reads were produced in an Illumina HiSeq 4000. Illumina adaptors and low-quality reads were removed for downstream analysis with *Trimommatic v.0.39* (134), parameters: *LEADING:3*; *TRAILING:3*; *SLIDINGWINDOW:4:15*; *HEADCROP:10*; *CROP:90*.

### TE annotation

For each sequenced genome, we developed a novel pipeline combining automatic and manual approaches to construct TE consensuses. First, we used *Earl Grey v.2.2* (135) to build de novo TE libraries with *RepeatModeler2 v.2.0.4* (136), which uses *RECON v.1.08* (137), and *RepeatScout v.1.0.6* (138). We used the following parameters: -r 32281 (NCBI taxon ID for *Drosophila*), -c yes (cluster redundant consensus), -m yes (remove consensus with less than 100 nt), and -e yes (enable *HELIANO v1.2.1* (139) to identify Helitrons). In short, EarlGrey automatically retrieves TE consensus, and performs an extension of each sequence using a method named “Blast, Extract, Extend” (BEE) (140). Such method extracts the sequences of the 20 full-length (or longest) TE insertions for each consensus, and then it retrieves the 1 kb flanking regions from them. Posteriorly, insertions altogether with flanking regions are aligned with *MAFFT v.7.487* (141), and the alignment is trimmed with *trimAl v.1.4* (142), parameters *–gt 0.6* and *–cons 60*. Subsequently, the alignment is analyzed to check whether the flanking regions from the full-length insertions have high quality, meaning that they are part of the TE rather than genomic regions. Finally, the consensus sequence for each TE family is updated with *EMBOSS v.6.6.0.0* (143). The BEE method is repeated up to 10 times for each consensus.

Once the extended TE libraries were obtained for the six genomes with EarlGrey, we implemented our pipeline into eight steps. Firstly, we removed all sequences other than TEs from EarlGrey’s library: satellites, low complexity repeats, tRNA, and rRNA. At step 2, we checked the presence of redundant consensus respecting the 80-80 (35) rule with all-vs-all blastn (-qcov_hsp_perc 80 and -perc_identity 80). Surprisingly, ∼45% of the consensuses were duplicated. Some of them were identical, or nearly identical, but in opposite strands. We selected the longest consensus from the ones considered redundant. At step 3, we used Tandem Repeat Finder (144) to identify consensuses that were had over 50% annotated as tandem repeats. Consensus falling in this cutoff were removed for being indistinguishable from a tandem repeat. At step 4, we removed all consensus with high similarity with *D. mojavensis wrigleyi* CDSs using blastn, parameters *-qcov_hsp_perc 80 and - perc_identity 80*. TE consensuses that had passed our filtering thresholds were then submitted to the classification analysis. At step 5 of the pipeline, TE consensus will be classified at the family level based on similarity analysis of non-reference consensus with a database provided by the user (https://github.com/OliveiraDS-hub/Pipelines-Cactophilic-Drosophila-Species). In our case, we used all consensuses from *Drosophila* species available on Dfam v3.7 (145). Due to the “curated” Dfam library containing only *D. melanogaster* consensus, we used the “uncurated” library from Dfam. We extracted *Drosophila* TEs with famdb.py v.0.4.3, parameters -i Dfam.h5 families --include-class-in-name -f fasta_name -ad ‘Diptera’. Then, we filtered out non-Drosophila sequences and “Unknown” TEs with the Unix command line: grep -i ‘drosophila’ file.fa | grep -v -i ‘unknown’, obtaining 49,916 out of 344,562 consensus. The pipeline firstly classifies at the family level by doing a blastn (- qcov_hsp_perc 80 and -perc_identity 80) between the *Drosophila* Dfam consensus and the provided de novo consensus. Approximately 50% of the identified consensus were classified on the family level based on this blastn. The consensus without results from this stringent blastn and smaller than 200 nt were filtered out. Due to the stringency of the matches with blastn, we also masked the remaining unclassified consensus with the *Drosophila* Dfam consensus with RepeatMasker.v4.1.8 (146). Based on RepeatMasker output, we computed the length of all matches from a given family in the unclassified consensus, and how much it has been covered based on the unclassified length. If the coverage is > 80% of the total length, we assume that the unclassified consensus has diverged from that one of the database, but it still classified as the same matched family. This step classified ∼25% of our consensuses. Finally, the remaining unclassified sequences that could not be classified at the family level are classified at the class/order level. We used a cutoff of 50% of covered matches in the unknown consensus to classify their classes. As the family name, we substituted the pattern from RepeatModeler2 as (i.e. rnd-1_family-1) to a pattern representing: “species preffix”, “author initials”, “homology percentage” (i.e. Dari_DSO_86.0275p#DNA/TcMar-Tc1). In this example, “Dari” stands for *D. arizonae*, “DSO” is the author’s initials, and “86.0275p” represents the percentage of the total length that this consensus has been matched with a DNA/TcMar-Tc1. The remaining putative TE consensuses with less than 50% of coverage with known *Drosophila* TEs were removed. At the end of this step, we end up with polished TE consensus.

In step 6, we checked the strandness of TE consensus sequences based on stranded RNA-seq data. During the clustering of similar sequences and removal of redundancy, we noticed many consensuses with nearly identical sequences, but in different senses. To correct TE strandness, we used our stranded RNA-seq data. We annotated TE insertions in the genome with *RepeatMasker v.4.1.4* (122), parameters *-cutoff 250, -s*, and *-norna*. Then, we mapped the reads from a mixed library from head and larvae tissues with STAR v2.7.11b (147). To be certain about the strandness origin of the reads, we removed all insertions that have any overlap with other TEs and/or exons. Then, for each consensus, the pipeline identifies the three copies in the genome with higher expression. Based on these copies, the number of uniquely mapped reads to the same strand of the copy is quantified. If > 50% of the reads align to the same strand of the copies, we assume the consensus strandness was right. Otherwise, the pipeline converts the consensus sequence to its reverse complement. This step will add a strandness tag to the consensus ID. For instance, if 75% of the reads support forward strand, a tag “75st” will be added to the family name. This is useful to provide a metric of certainty about the strandness of each consensus. Such information can be valuable when working with piRNAs, where alignment strandness is crucial to interpret the results. Then, we used *RepeatCraft* (148) with *LTR Finder v.1.2* results to merge LTRs and their respective internal regions. Subsequently, we removed all TE insertions shorter than 80 nucleotides, following the classification rule 80-80-80 (35). Finally, we performed a last filtering step in the TE annotation to remove insertions composed of simple sequence repeats. We used *TRF v.4.09* (144) in all TE insertions from each genome, with parameters *2, 5, 6, 75, 20, 50, 500, -m, -h*, to remove TEs with masked repeats over more than 50% of their sequences. This method has been previously used to remove TE copies that are indistinguishable from simple repeats (24,149).

### TE enrichment at regulatory regions

To identify TE insertions within and nearby genes for each genome, we created *bed* files with the genomic positions of all insertions. Then, we created *bed* files for the gene regions corresponding to the 2 Kb upstream of the transcription start-site (TSS). To test whether the prevalence of TEs in each gene region was significantly higher than expected (enriched) compared to random genomic distributions of TEs in the same region of other genes, we performed permutation tests with the R package *regioneR* (150), parameter *randomization = resampleRegions*, with 2,000 random samples. For each gene family with *n* genes, the number of genes with TEs was compared to other random *n* genes 2,000 times. This analysis allowed us to infer whether there are HLAU gene families with the observed frequency of genes with TEs at the promoter region higher than expected compared to the rest of the genome. We also selected zinc-finger proteins and serine/threonine kinases as two non-HLAU gene families to compare our findings. Taking into account that TEs might harbor regulatory and/or coding motifs in both strands (62), our analysis considered TEs within the investigated gene regions regardless of the strand.

### Identification of TE-rich genomic regions

We split the genomes into bins of 200 kb, generating a bed file with such intervals across the genomes. Then, we used the function *intersect* from *bedtools v.2.30.0* (151) to identify the number of TE insertions located in each genomic bin. Posteriorly, we merged all overlapped TE insertions into single ranges. These ranges were summed to obtain the total TE content in bp for each 200kb bin. By plotting the TE content of each genome, we noticed that bins with more than 50kb of TE content were outliers of the normal distribution. Therefore, we defined TE-rich regions as all bins with > 50kb of TE content, which represents 25% of the bin size. We used the same function from *bedtools* to identify whether OR genes were present in the TE-rich regions, or the TE-poor (bins with less than 50kb of TE content).

### Transcription factor binding sites on TE insertions

We assessed the identification of transcription factor binding sites (TFBSs) with *FIMO* (152), from the *MEME v. 5.0.5* suite tools (153). We extracted the sequences of all TE insertions located 2kb upstream of ORs with *bedtools v. 2.30.0*, function *getfasta*, using parameter *-s* (151). We obtained the TFBS motifs described for *D. melanogaster* from Jaspar database (154), for the transcription factors (TFs) acj6, zf30c, fer1, onecut, and xbp1, which have been demonstrated to activate ORs (43). To compare our findings with OR genes, we performed the same analysis with Serine/Threonine Kinases and zinc-finger as non-HLAU gene families. Only TFBSs with p-value < 0.01 were considered. The enrichment of OR TFBSs in OR genes compared to non-HLAU gene families was assessed with the four frequencies: 1) ORs with TFBSs; 2) ORs without TFBSs; 3) non-HLAU with TFBSs; and 4) non-HLAU without TFBSs.

### Differential expression of genes

We aligned the RNA-seq reads with *STAR v.2.7.3* (147), using the assembled genomes and annotations of their respective species. The alignments were used to create the count matrix reporting the expression of each gene, with *htseq v.0.13.5* (155), parameters *–order pos*; *-s reverse*; *–i gene*; *-t exon*; the latter indicates that the count for each gene corresponds to the sum of all reads aligned into exons (CDSs and UTRs). We merged the counts from species of the *buzzatii* and *mojavensis* clusters separately, maintaining only ortholog genes that have at least 90% of the length of the annotated gene in the reference genome. This threshold allowed us to avoid misinterpretation of the expression level by comparing genes with different lengths. Then, we performed independent statistical analysis for species of the *buzzatii* and *mojavensis* clusters with *DEseq2* v.1.28.1 (156), using a single-factor experimental design, which considers only the factor that different species may have differentially expressed genes. Due to the phylogenetic distance between species of the *buzzatii* and *mojavensis* clusters, we performed the differential expression analysis separately for each group. We only considered genes with adjusted *p-value* < 0.05 and |*log2 fold-change*| > 1 as differentially expressed. To test for association between TE enrichment and differential expression, we selected HLAU genes with the mean of normalized counts between replicates >= 100. Then, we build the contingency table to apply the *X*^2^ independence test, splitting them into four categories: differentially expressed, non-differentially expressed, with TEs at the promoter region, and without TEs at the promoter region. We carried out the test for all pairwise analyses between cactophilic species with preference for *Opuntia sp* and columnar cacti. *X*^2^ with p-value < 0.05 was considered significant. The proportional differences between localization, acceptance, and host use genes compared to non-HLAU genes (Ser/Thr Kinases and zinc-finger gene families) were assessed with Fisher’s exact test, with a contingency table containing the average number of genes per group: 1) non-HLAU differentially expressed genes; 2) non-HLAU non-differentially expressed genes; 3) HLAU differentially expressed genes; and 4) HLAU non-differentially expressed genes. *Fisher’s* Exact test < 0.05 was considered significant.

### Identification of gene-TE chimeric transcripts

We detected gene-TE chimeric transcripts with *ChimeraTE v.2.0.0* Mode 1 (24). Each species was analyzed using its respective genome, gene annotation, and TE annotation. Shortly, *ChimeraTE* identifies chimeras based on paired-end reads spanning from exons to TE sequences. Based on the TE position compared to the gene (upstream, exon, intron, downstream), chimeric transcripts are classified into categories. We have used default parameters, except *--replicate 3*, which considers true chimeras only those identified in the three RNA-seq replicates, and *–coverage 10* to obtain chimeras with at least 10 chimeric reads on average between replicates. To calculate the contribution of the chimeric transcript expression relative to total gene expression, we developed a new module for ChimeraTE. This module can be used downstream to ChimeraTE mode 1 analysis. In the first step, all RNA-seq replicates will be merged into a single fastq library. Then, the merged reads are aligned to the genome with STAR (147). Next, Stringtie2 (157) is used to assess transcript expression, including both reference and non-reference isoforms. All TEs generating chimeras are intersected with exons predicted by Stringtie with pyranges (158). Only overlaps with 80% of the TE within the exon are considered. Finally, the expression of the TE-containing isoforms is compared to the total gene expression. In our analysis, we removed all chimeric transcripts for which Stringtie was not able to build the chimeric isoform. In addition, aiming to keep chimeras with putative function, we selected only the ones with contribution >= 25%. This analysis can be reproduced with any output from ChimeraTE with the code “mode1_contrib_chimeras.py” (https://github.com/OliveiraDS-hub/ChimeraTE/blob/main/scripts/mode1_contrib_chimeras.py).

### Data access

The quality control data from ONT and RNA-seq data, the genome assemblies, and their respective gene and TE annotation (libraries and gtf files) are available in the Zenodo repository: https://zenodo.org/records/16501018. The codes to reproduce differential expression analysis, positive selection tests, gene annotation, permutation tests, TE annotation, and TFBS analysis are available on the GitHub repository: https://github.com/OliveiraDS-hub/Pipelines-Cactophilic-Drosophila-Species.

## Acknowledgements

We thank Séverine Chambeyron for useful discussions and advice. This work was performed using the computing facilities of the CC LBBE/PRABI. This study was funded by Brazilian agencies FAPESP-The São Paulo Research Foundation [grant number 2020/06238-2] and from the National Council for Scientific and Technological Development (308020/2021-9) to C. M. A. C; the Eiffel Fellowship Campus France [P769649C] and CAPES – PrInt [88887.716810/2022-00] to D. S. O.; by Ministerio de Ciencia e Innovación (Spain) [grant number PID2021-127107NB-I00], and Generalitat de Catalunya (Spain) [grant number 2021 SGR 00526] to M. P. G. G.; and by Udl-FAPESP ANR-IDEX-0005 to C. V. This study was financed in part by the Coordenação de Aperfeiçoamento de Pessoal de Nível Superior – Brasil (CAPES) – Finance Code 001 to DSO.

## Authors’ contributions

A. L., A. B, M. P. G. G, performed experiments; F.S., Performed genome assemblies; W. V. B. N. performed phylogenetic analysis; D. S. O performed computational analysis related to genomics, transcriptomics, and statistical tests; D. S. O. wrote and edited the paper; C. V. and C. M. A. C. designed the study, and reviewed the manuscript. All authors read, corrected, and approved the final manuscript.

