## Supplementary figures and images for "Transposable elements contribute to the evolution of host shift-related genes in cactophilic *Drosophila* species"

### Supplemental Fig. S1

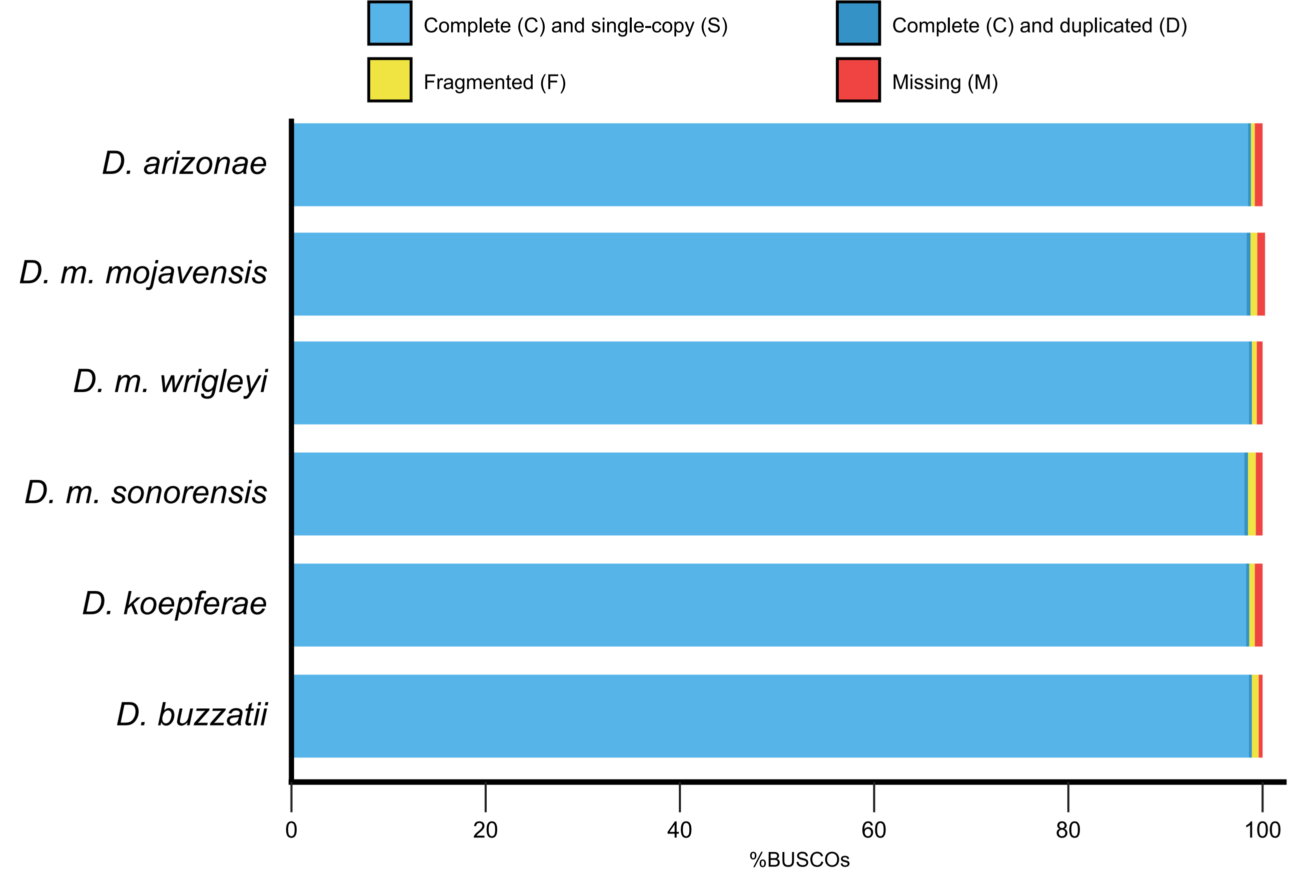

### Supplemental Fig. S2

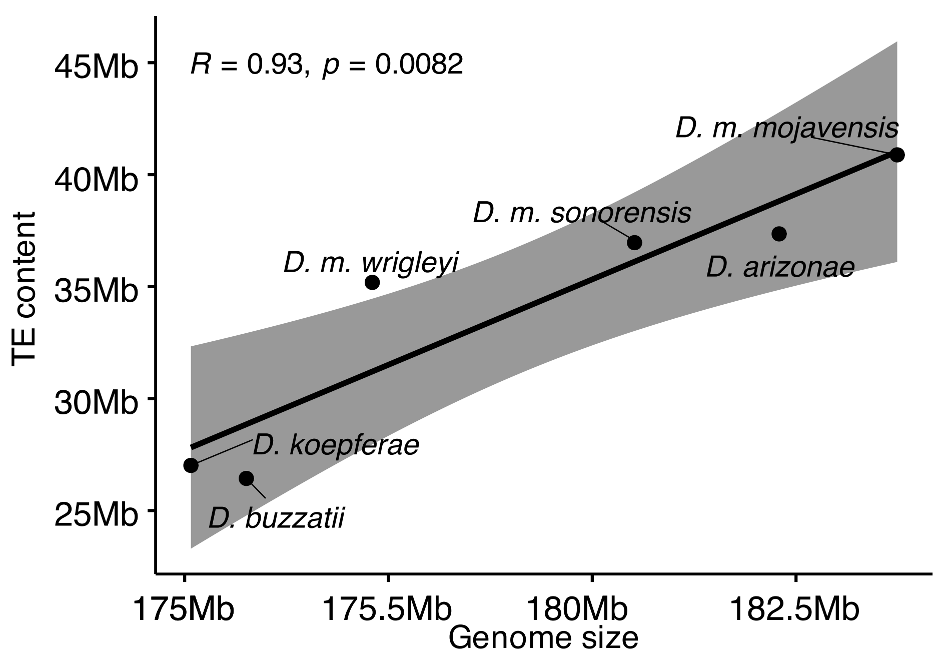

### Supplemental Fig. S3

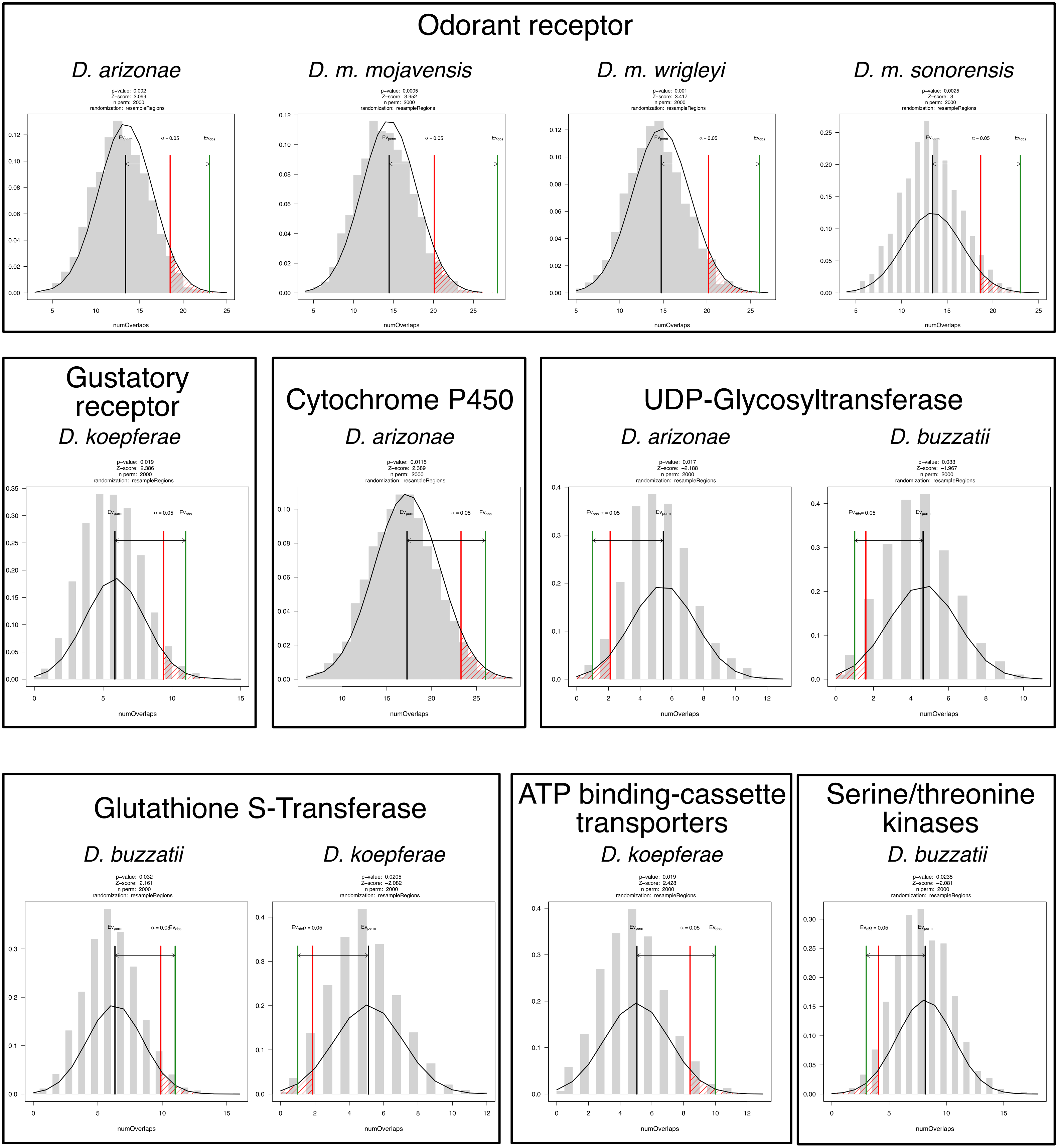

### Supplemental Fig. S4

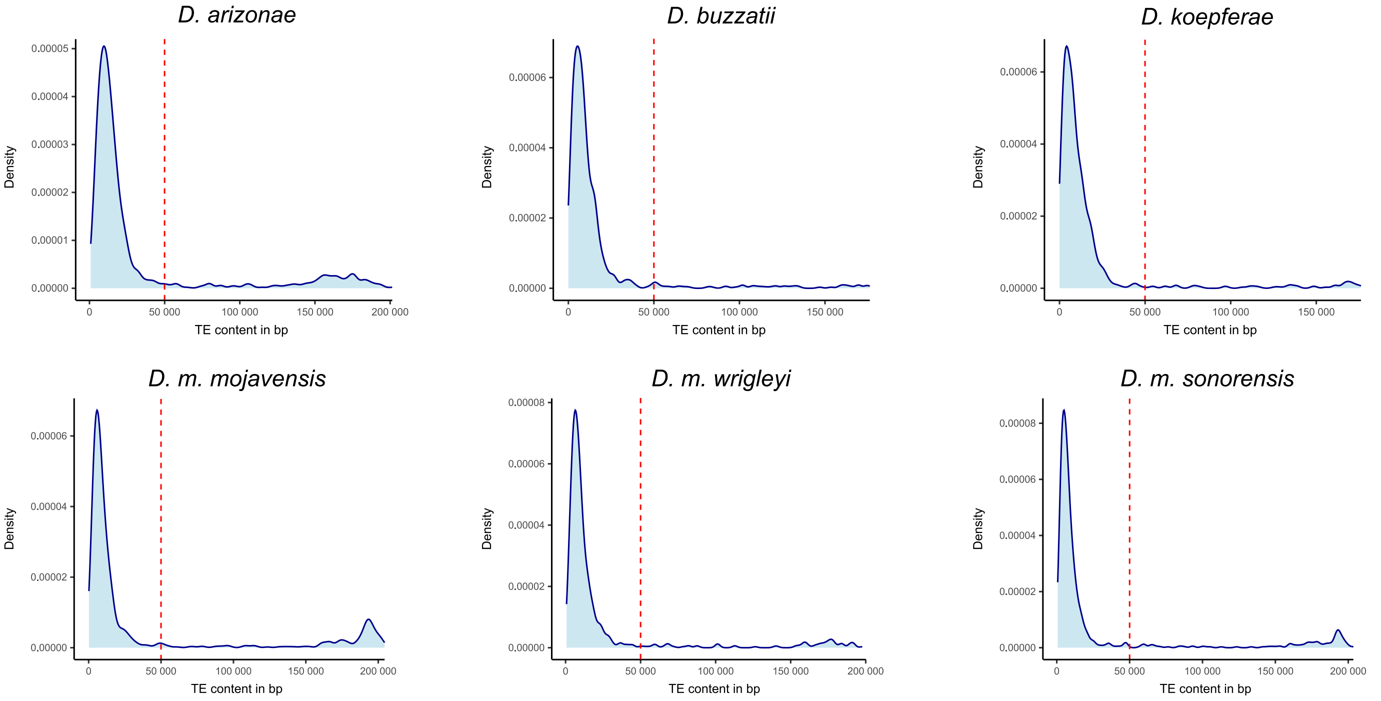

### Supplemental Fig. S5

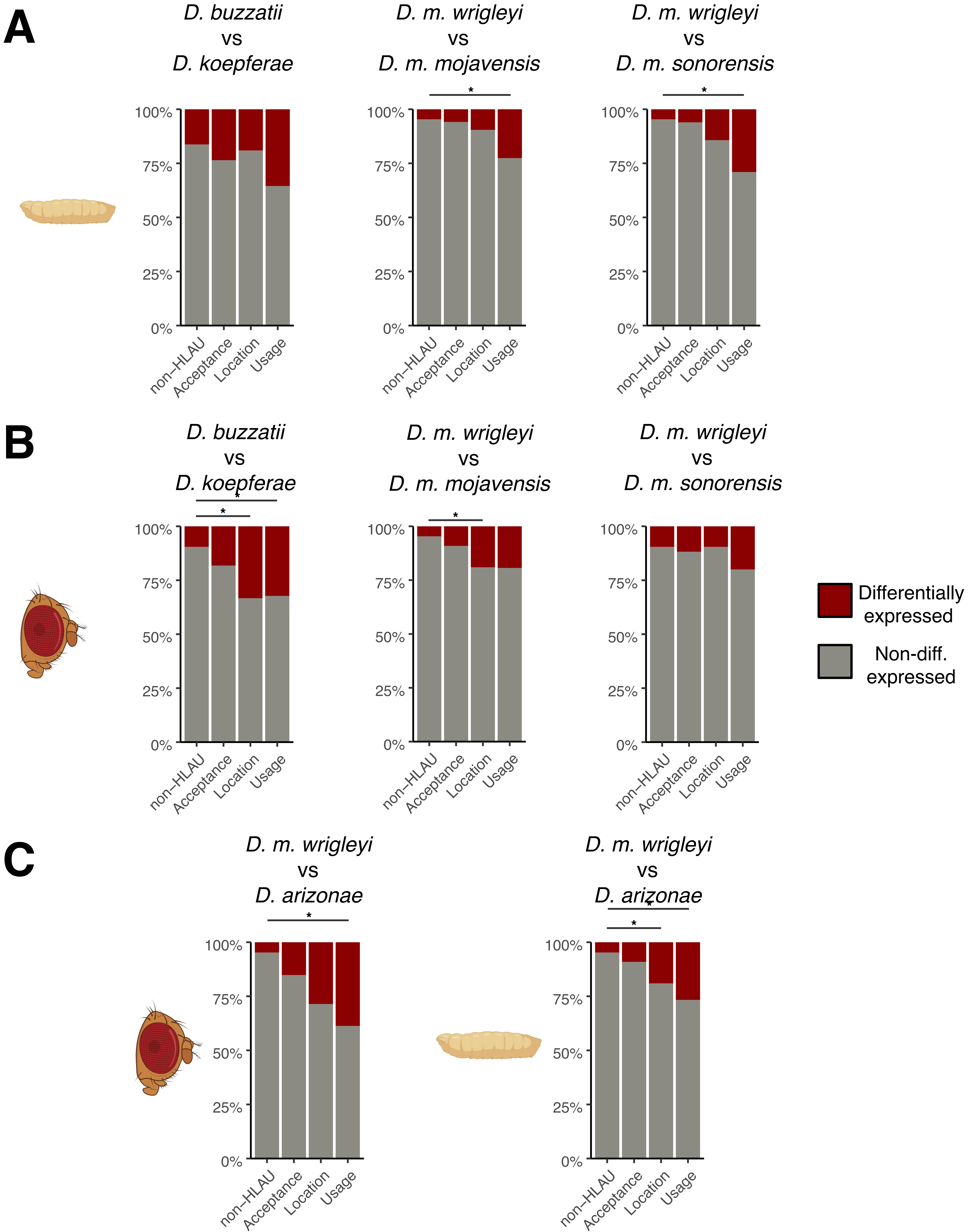

### Supplemental Fig. S6

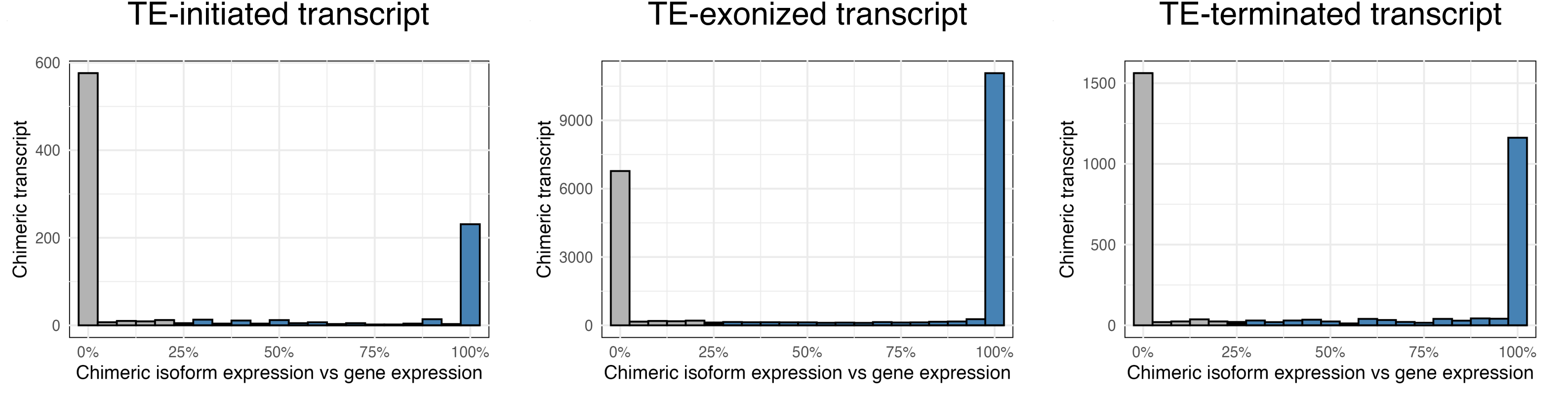
